# An *Aspergillus* lncRNA atlas reveals a novel modulator of aflatoxin biosynthesis

**DOI:** 10.64898/2026.05.29.728754

**Authors:** M. Laura Fabre, Jin Woo Bok, Benjamin J. Haefner, Nancy P. Keller, Amelia E. Barber

## Abstract

Long non-coding RNAs (lncRNAs) represent a significant fraction of the transcriptome and are vital regulators in eukaryotes, yet their functional landscape in fungi remains largely undefined. Here we present a transcriptomic atlas of 8,553 lncRNAs across four evolutionarily and biotechnologically diverse *Aspergillus* species. We observed marked diversity in the long non-coding repertoire between species, with only 39% shared between at least two species. However, conserved lncRNAs are expressed at significantly higher levels and across more conditions than species-specific ones, revealing a functional hierarchy within the non-coding transcriptome. We also identify subtelomeres and transposable elements as hotspots for lncRNA innovation in *Aspergillus*, albeit at the cost of reduced expression. Integrating genomic and transcriptomic data, we show that lncRNAs predominantly exhibit distance-dependent *cis-*coexpression with neighboring genes and, through guilt-by-association, we identify their participation in global modules governing primary and secondary metabolism. This framework enabled the discovery of *aflalinc,* a species- and condition-specific lncRNA in *A. flavus* that plays a role in regulating the aflatoxin biosynthetic gene cluster. Crucially, deletion of this lncRNA severely reduced the production of aflatoxin *B₁*, a highly potent agent of acute toxicity and driver of hepatocarcinoma, without affecting growth, conidiation, or sclerotia formation. In addition, examination of an independent dataset confirms the strength of the co-expression networks and uncovers stage-specific functions for lincRNAs in the maturation of *A. fumigatus* biofilms. Together, these findings establish lncRNAs as integral components of fungal biology and demonstrate that they can modulate mycotoxin production and biofilm formation, with broad implications for controlling fungal threats to food safety and human health.

## Introduction

Long non-coding RNAs (lncRNAs) are vital regulators of gene expression across eukaryotes, acting as molecular scaffolds, decoys, and guides that shape chromatin architecture and transcriptional output [1,2]. These non-coding transcripts can occupy a significant fraction of the transcriptome, where on the upper end they outnumber protein-coding genes by a ratio of 2:1 in the human genome [3]. Despite this and their established roles in animals and plants, the landscape and functions of lncRNAs in fungi remain largely unexplored [4]. However, recent work has begun to establish important roles for fungal lncRNAs in virulence and secondary metabolism. In the emerging human pathogen *Candida auris*, the lncRNA *DINOR* modulates virulence via controlling stress responses and modulation of the DNA damage checkpoint kinase Rad53 and TOR signaling [5]. In *Cryptococcus neoformans*, *RZE1* regulates the yeast-to-hypha transition, a key virulence determinant, by controlling the transcription factor *Znf2* [6]. In the filamentous plant pathogen *Fusarium graminearum*, the lncRNA *RNA5P* acts as a *cis*-acting inhibitor of TRI5 expression, thereby suppressing synthesis of the mycotoxin deoxynivalenol [7]. Recent studies have also begun to highlight the functional relevance of lncRNAs in *Aspergillus*, including work in *A. fumigatus* showing that the lncRNA *afu-254* modulates stress responses and virulence, and additional evidence that other lncRNAs contribute to antifungal sensitivity [8,9]. While condition-specific studies have begun to reveal individual lncRNA functions, a systematic, cross-species understanding of how these transcripts differ across *Aspergillus* species has been lacking.

LncRNAs arise through diverse evolutionary mechanisms, including the domestication of transposable elements (TEs), pseudogenization of protein-coding genes, and *de novo* emergence from previously non-functional genomic regions [10–12]. Work in metazoa has highlighted that transposable element sequence is overrepresented within lncRNAs. This is believed to be due to the benefits provided by exonic TE sequences being incorporated as functional domains within lncRNAs. Called the RIDL hypothesis, standing for Repeat Insertion of Domains of LncRNAs, TE domains are repurposed by lncRNAs as the building blocks that provide the “pre-built” structural and sequence features that allow them to interact with and regulate other cellular targets [13]. In vertebrates, 66-75% of mature lncRNAs contain sequence from TEs [14], and similarly high proportions have been observed in plants, with 65% of maize lncRNAs estimated to be TE-derived [15]. Beyond mere structural integration, TE-derived domains have been functionally validated as essential regulatory modules [16,17]. However, the extent to which TEs have shaped the generation and evolution of lncRNAs in fungi, including in *Aspergillus*, has not been explored.

The genus *Aspergillus* is particularly well-suited for investigating the breadth and functional landscape of lncRNAs across a fungal taxon due to its diversity and human relevance*. Aspergillus fumigatus* is the leading cause of invasive aspergillosis, a devastating disease with a crude mortality of 1.8 million deaths per year [18]. *Aspergillus flavus* represents a major threat to food safety through contamination of crops with aflatoxin *B₁*, a group 1A carcinogen [19,20]. *Aspergillus niger* is an industrial model organism for organic acid and enzyme production [21], and *Aspergillus nidulans* serves as a foundational genetic and cell biology model [22]. Although they all belong to the genus *Aspergillus*, there is significant evolutionary divergence between the four species. The last shared ancestor between *A. fumigatus* and the other three species was an estimated 77 million years ago, during the upper cretaceous period [23]. Even the most closely related pairing of the four species, *A. nidulans* and *A. niger*, split at estimated 62 million years ago. While the clinical and economic importance of these four species is clear, the lncRNA landscape and its roles in coordinating key fungal traits, including virulence, secondary metabolism and industrial performance, remain poorly defined.

A hallmark of fungi, including *Aspergillus* spp, is their capacity to produce a vast array of secondary metabolites. These bioactive compounds mediate ecological interactions but also act as toxins to plants and humans [24–26]. Secondary metabolites are synthesized by physically clustered genes known as biosynthetic gene clusters (BGCs), which are often co-regulated in response to environmental cues [25]. Studies of BGC regulation have traditionally focused on cluster-localized transcription factors and chromatin modifiers [27,28], but the increased accessibility of RNA sequencing has facilitated global co-expression network approaches with the ability to discover distal, *trans*-acting regulators [29,30]. We posit that lncRNAs are integral players in this space, and that systems biology approaches can be extended to identify lncRNAs as functional partners that influence BGC expression.

Building on these established roles in other eukaryotes, we hypothesized that lncRNAs serve as active and abundant components of *Aspergillus* transcriptomes, and that the specific roles they play can be systematically uncovered through integrative network analysis and experimental validation. To test this, we integrated 993 RNA-seq samples across four species, reconstructed co-expression networks for the 8,553 lncRNAs discovered to identify their putative functional associations, and validated our predictions through two independent approaches: first, by experimentally disrupting *aflalinc*, an lncRNA co-expressed with the aflatoxin BGC in *A. flavus* and whose deletion resulted in significantly reduced aflatoxin production; and second by assessing module preservation and temporal dynamics using an independent *A. fumigatus* biofilm dataset, which revealed that lncRNAs exhibit stage-specific patterns beyond their protein partners. We also uncover a significant interplay between transposable elements and lncRNAs and strong distance-dependent *cis*-coexpression with neighboring genes, as well as systemic participation in global gene expression modules. Together, these findings provide a foundation for exploring the non-coding dimension of the fungal kingdom, with implications for controlling fungal pathogens and optimizing industrial strains.

## Results

### lncRNAs represent up to 18% of the total predicted gene content in *Aspergillus* spp

LncRNAs are increasingly recognized for their major contributions to gene regulation in animals and plants, but their landscape and potential contributions in fungi remains largely unexplored. To address this gap, we built a comprehensive catalog of lncRNAs across four evolutionarily and biotechnologically diverse *Aspergillus* species: *A. fumigatus, A. nidulans, A. niger and, A. flavus*. Using 993 publicly available RNA-seq samples (Suppl. Table S1) and a state-of-the-art computational pipeline (Fig. 1A), we identified 8,553 long non-coding genes (*A. fumigatus*: 1,165; *A. nidulans*: 2,291; *A. niger*: 2,356; *A. flavus*: 2,741), which increased the total gene count by approximately 10-20% for each species (Fig. 1B, Suppl. Fig. S1A, Suppl. Table S2). The number of lncRNAs identified was broadly proportional to each species’ genome size (Pearson correlation coefficient = 0.82; p = 0.18) and was not dependent on the number of RNAseq samples used. *A. fumigatus* had the largest number of transcriptomes available (n=553). When these were downsampled to match the number of RNAseq samples for the smallest dataset (*A. nidulans,* n = 120), the number of lncRNAs predicted was similar for both the full and reduced datasets (n = 1,165 and n = 1,203, respectively).

**Figure 1.**
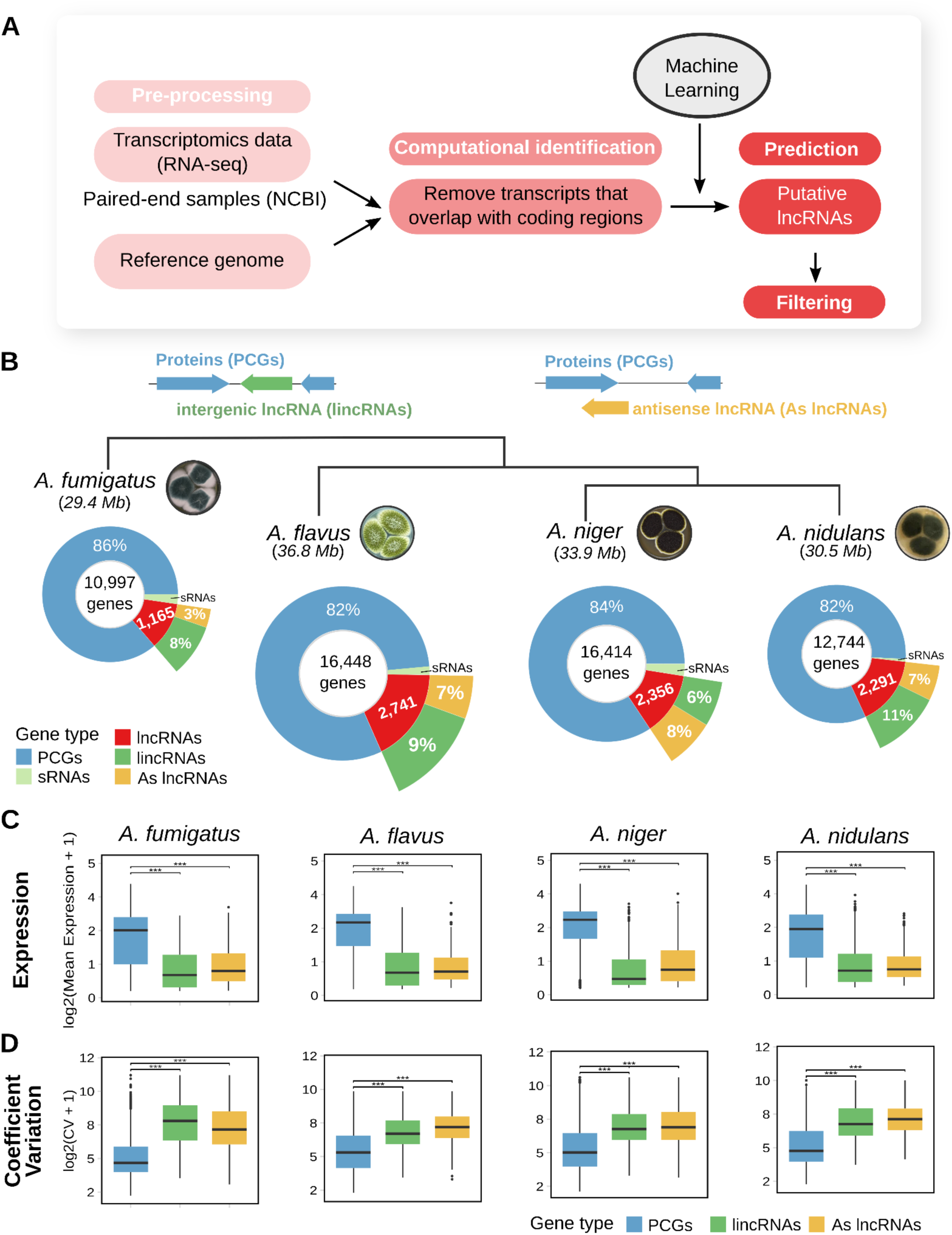
Computational identification, genomic distribution, and expression characteristics of lncRNAs in four model *Aspergillus* species. (A) Overview of the lncRNA discovery workflow. Public RNA-seq datasets and reference genome assemblies were used for transcript assembly (Suppl. Table S1). Existing annotated genes were excluded and machine learning-based classification was applied to predict putative lncRNAs, followed by filtering. (B) Schematic representation of antisense (As lncRNAs) and intergenic lncRNAs (lincRNAs) (top). Genomic composition of *A. fumigatus, A. flavus, A. niger,* and *A. nidulans*, showing proportions of protein-coding genes (PCGs), small RNAs (sRNAs) as present in the existing reference genome annotation, and the novel lncRNAs predicted in this study (Suppl. Table S2). The total number of genes (lncRNAs and PCGs) for each species is indicated at the center of each plot (bottom). Fungal plate images were adapted from Wikipedia, CC BY-SA 2.5 (*A. fumigatus*) and CC BY-NC 4.0. (C-D) Distribution of expression levels (C), and coefficient of variability (CV) (D) for protein-encoding transcripts, lincRNAs, and As lncRNAs. Expression values are log2-transformed for clarity. Statistical significance assessed using the Wilcoxon rank sum test with Bonferroni correction; *** p < 0.001, ** p < 0.01, * p < 0.05.

Due to the ambiguity associated with studying lncRNAs that are sense to protein-coding genes (PCGs), we focused on those that are antisense (As lncRNAs), located on the opposite strand of a protein-coding gene, and those that are intergenic (lincRNAs), located between protein-coding genes. LincRNAs were the dominant class after protein-coding genes in *A. fumigatus* (8%), *A. nidulans* (11%), and *A. flavus* (9% of total genes), while antisense lncRNAs comprised a smaller fraction (3-7%) in these species. *A. niger* was the exception, with slightly more As lncRNAs than intergenic lincRNAs (995 vs. 902 genes), a pattern likely resultant from its less contiguous scaffold-level genome assembly, making it more challenging to identify transcripts in intergenic regions. Structurally, lncRNAs exhibited a median of two exons per transcript across all species (Suppl. Fig. S1B), substantially fewer than protein-coding genes, which showed a median of three exons. This simpler exon structure is consistent with the reduced splicing complexity often observed for lncRNAs in other eukaryotes [31] (Suppl. Fig. S1B).

Having established a comprehensive catalog, we next investigated the lncRNAs for features that differentiated them from protein-coding genes. In vertebrates and plants, lncRNAs are typically longer and expressed at lower levels than protein-coding genes and exhibit higher expression variability across conditions and tissues [2,32,33]. Our analysis revealed that *Aspergillus* lncRNAs are generally shorter than PCGs, a pattern also observed in *Neurospora crassa* and other fungi [34–36] (Suppl. Fig. 1C). They also exhibited lower expression levels and significantly higher expression variability across the diverse conditions sampled, as measured by coefficient of variation (Fig. 1C). These features, particularly their lower and more variable expression, are consistent with the model that lncRNAs in *Aspergillus* act as context-dependent regulators rather than constitutively expressed components of cellular function, a *modus operandi* also observed in other eukaryotic organisms [2,4]. Taken together, we find that lncRNAs represent a notable, but previously unappreciated portion of the *Aspergillus* genome and that their expression is much more condition-dependent than protein-coding genes.

### 488 lincRNA families are conserved between at least species and are expressed at higher levels

We next sought to quantify the degree to which our newly identified *Aspergillus* lncRNA catalog was conserved among the four species. While PCGs are subject to stringent constraints to maintain open reading frames, lncRNAs exhibit high evolutionary plasticity at the primary sequence level, which allows them to evolve much more rapidly compared to the proteome [14,37]. To avoid the confounding evolutionary constraints imposed upon As lncRNAs, we focused our conservation analysis on lincRNAs. Due to the high degree of sequence plasticity in lnRNAs, we used a synteny-based approach as in Pegueroles *et al*., 2019 to identify putatively orthologous lincRNAs based on their location between conserved flanking protein coding genes (Fig. 2A)[38]. This approach identified 488 lincRNA families that were syntenically conserved across at least two species. In line with the relaxed evolutionary constraints on lncRNAs and their typical roles in regulation rather than essential cellular functions, species-specific lincRNAs outnumbered conserved lincRNAs in all species but *A. niger* at a ratio of 1.24 to 2.26x (Fig. 2B, Suppl. Table S4). In *A. niger,* the higher fragmentation of its genome likely obscured the detection of some conserved lincRNA, as the necessary syntenic flanking anchors are more likely to have been missing. The degree of lincRNA family conservation was also reflected by phylogenetic distance. *A. fumigatus* is the most phylogenetically divergent species in the dataset and contained the fewest number of the conserved families (n=167; Fig. 2C, Suppl. Table S4), while *A. flavus* harbored the highest number (n=398). However, 34% of families were conserved between *A. fumigatus* and at least one other species, illustrating an appreciable ancestral lincRNA backbone.

**Figure 2.**
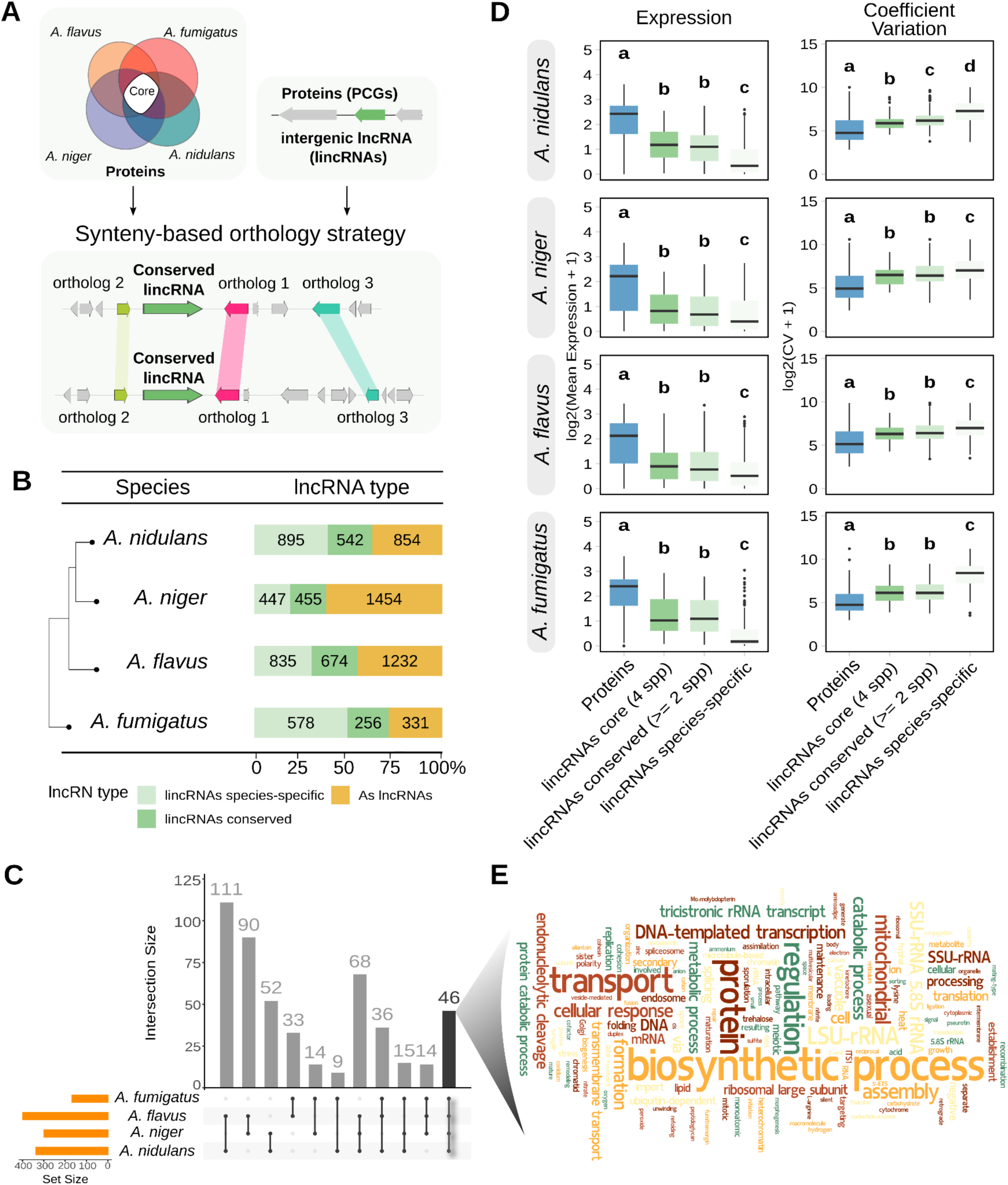
Identification of conserved intergenic lncRNAs in *Aspergillus* reveals their higher expression and lower variation relative to species-specific lincRNAs. (A) Schematic overview of the synteny-based strategy used to identify conserved lincRNAs. lincRNAs were considered conserved if they were flanked by at least one single-copy protein-coding orthologous gene in the up- and downstream regions and if a third orthologous protein-coding gene was shared in the three protein coding genes up- and downstream from the lincRNA. (B) Bar plot showing the percent and number of antisense lncRNAs, conserved (present in ≥2) lincRNAs, and species-specific lincRNAs. (C) UpSet plot illustrating the overlap of conserved lincRNAs families among the species. (D) Comparison of expression (TPM) and coefficient of variability (CV) between protein-coding genes, conserved lincRNAs, and species-specific lincRNAs across all species. Statistical significance was assessed using the Wilcoxon rank sum test with Benjamini-Hochberg correction. Letter groupings indicate statistically distinct groups at p < 0.05, where groups sharing a letter are not significantly different. (E) Word cloud of GO term names and definitions from WGCNA co-expression modules containing the 46 core lincRNAs families conserved across all four *Aspergillus* species. Word size reflects the relative frequency of each biological process term across the modules harboring these core lincRNAs.

To explore the relationship between evolutionary conservation and strength of transcriptional activity, we compared the expression of lincRNAs conserved between at least two species and those that were species-specific (Fig. 2D). We hypothesized that more ancient lincRNAs that have persisted in the face of purifying selection are more likely to mediate central regulatory functions, while species-specific transcripts evolve rapidly to support niche adaptation. As a result, species-specific lincRNAs may therefore be expressed only under specific conditions and at overall lower levels, as they have had less evolutionary time to be integrated into regulatory circuits. Consistent with this, we discovered a clear transcriptional hierarchy: PCGs were the most highly expressed, followed by the conserved lincRNA, and species-specific lincRNAs exhibited the lowest expression levels (Wilcoxon rank sum test, p < 0.001; Fig. 2D). Species-specific lincRNAs also were the most variable in expression, as demonstrated by significantly higher coefficients of variation than both conserved lincRNAs and PCGs (Wilcoxon rank sum test, p < 0.001 for all species). This pattern supports the idea that these lowly expressed, species-specific transcripts are a nascent pool of regulatory material that is not as incorporated into cellular networks, but provides the plastic genetic material needed for species-specific adaptation and environmental responses. Notably, we identified a core set of 46 lincRNA families that were synteny-conserved across all four species (175 total transcripts and representing 9% of total lincRNA families) (Fig. 2C). The expression and condition-specificity of these core lincRNAs were not significantly different to lincRNAs conserved across only two to three species (Fig. 2D). However, given the faster evolution of non-coding genes, the fact that these lncRNAs have been conserved across all four species for more than 75 million years of evolution, suggests that they are involved in fundamental processes. In support of this, analysis of the Gene Ontology (GO) annotations of the protein-coding genes that were strongly co-expressed with core lincRNAs revealed that they are involved in central biological processes including myriad primary biosynthetic processes, transport, and translation (Fig. 2E). Altogether, evolutionary conservation analysis revealed that the largest fraction of lincRNAs is species-specific, in line with roles in adaptation to new environments, but conserved and core lincRNAs are more robustly expressed and implicated in central fungal biology.

### Transposable elements and subtelomeres as hotspots of lincRNA origin

Given the rapid evolution of lncRNAs in general and the high degree of species-specific non-coding transcripts observed, we next investigated whether their genomic distribution favored known regions of higher chromosomal instability. Subtelomeric regions are recognized hotspots of gene expansion, secondary metabolism, and pathogenicity in filamentous fungus [39–42]. Taking the terminal 10% of each from chromosome arm as the subtelomeric space, we detected a significant enrichment of lncRNAs (including both antisense and intergenic) in the subtelomeric regions of *A. fumigatus* (Chi-Square χ² = 192.46, p < 0.001) and *A. nidulans* (Chi-Square χ² = 29.33, p < 0.001) (Suppl. Fig. S1C; Suppl. Table S2). *A. flavus* showed no significant subtelomeric enrichment (Chi-Square χ² = 2.51, p = 0.112). *A. niger* was excluded from this analysis due to the absence of a chromosome-level genome assembly.

We hypothesized that the enrichment for lncRNAs in subtelomeric regions was driven by transposable elements, as we also identified a subtelomeric enrichment for transposable elements in all three species with chromosome-level assemblies (Pearson’s chi-squared test, all χ² > 10^4^, p < 0.001). To avoid the confounding effects of analyzing lncRNAs that overlap protein-coding sequences, we focused the next set of analyses on intergenic lncRNAs. To verify that none of the lincRNAs we identified were actually intact transposable elements, we first performed BLASTX searches against a curated transposon protein database (mycoMobilome) using relaxed parameters (e-value < 1e-5, coverage > 50%, identity > 30%). No significant matches to transposase or reverse transcriptase domains were detected, supporting that the lincRNA are *bona fide* non-coding transcripts and not simply misidentified, intact transposons. When we examined the proportion of lincRNAs that contained ≥10 bp of TE sequence, we found that *A. fumigatus* and *A. nidulans* exhibited the highest rates (44% and 34%, respectively), while only 21% of *A. niger* and 19% of *A. flavus* lincRNAs contained TE sequence (Fig. 3A, Suppl. Table S4). The integrated repeat landscape was dominated by DNA transposons and LTR retrotransposons across all four species (Fig. 3A). However, roughly half of the TE sequence contained within lincRNAs were labeled as unclassified, likely reflecting strong sequence decay post-integration.

**Figure 3.**
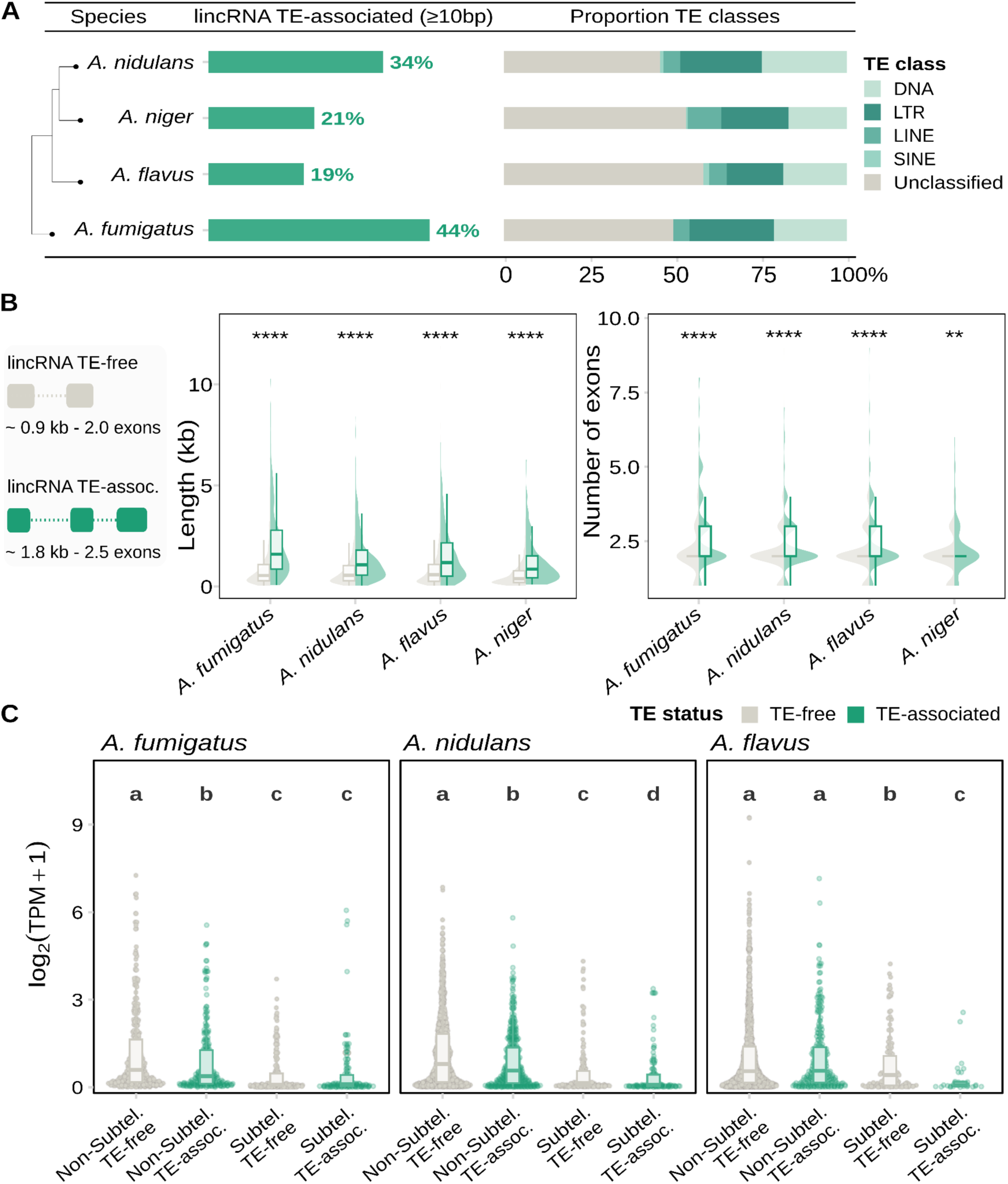
TE-associated structural and expression features of *Aspergillus* intergenic lncRNAs. (A) Bar plots represent the percentage of the lincRNAs repertoire containing exonic sequences that intersect with annotated class I and class II TEs. (B) Comparison of transcript length and exon number between TE-free (grey) and TE-associated lincRNAs (green). Boxplots show median and interquartile range; notches indicate 95% confidence intervals for the median. Significance for TE-free vs. TE-associated lncRNAs was assessed using Wilcoxon rank-sum test; *** p < 0.001, ** p < 0.01. (C) Influence of subtelomeric localization on lincRNA expression. Significance was assessed using Wilcoxon rank-sum tests with Benjamini-Hochberg correction. Letters indicate statistically distinct comparisons at p < 0.05, where groups sharing a letter are not significantly different.

Consistent with the RIDL hypothesis, where TE fragments provide functional building blocks to nascent lncRNAs, TE-associated lincRNAs were significantly longer across all species and exhibited a more complex transcript structure with higher exon counts (Wilcoxon rank sum test, p < 0.05 for all; Fig. 3B). TE-associated lincRNAs were roughly double the length of TE-free (0.7-2.4 kb vs. 0.4-1.2kb depending on the species) and had an average of 2.3-2.9 exons compared to 2-2.2 exons for lincRNAs where no TE sequence was detected. When we examined the percentage of TE sequence that made up each lincRNA, we found that subtelomeric lincRNAs contained significantly more TE sequence than those in other genomic regions, with no enrichment for any specific TE class (Wilcoxon rank sum test, p < 0.0001; Suppl. Fig. S2A, B). This was most strongly demonstrated in *A. fumigatus*, where subtelomeric lincRNAs were composed of a median of 21% TE sequence, compared to 8% for other genomic regions. Given the enrichment in subtelomeric regions and the high proportion of lincRNAs that contained TE sequence, we next wanted to look how these two facets might affect lincRNA expression, given that TEs are typically silenced. Interestingly, the combination of subtelomeric location and TE-association resulted in the lowest expression among all lincRNAs (Fig. 3C). We posit that this may be a residual consequence of the heterochromatin-mediated silencing that typically occurs following a novel TE integration, but may also represent a more intentional balance between allowing the innovation of lncRNA novelty and mitigating potential fitness costs. Taken together, we find that up to 43% of lincRNAs contain TE sequence in *Aspergillus* spp. and this is even more amplified for those located in the subtelomeric regions.

### LincRNAs display distance-dependent *cis*-associations with protein-coding genes

As lncRNAs largely function in a regulatory context, mediating the expression of protein coding genes rather than directly performing a biochemical function, we next sought to better understand how they regulate gene expression in *Aspergillus*. We first tested for local, *cis-*acting interactions by examining the correlation of lncRNA expression with neighboring PCGs within 15 kb. Across all species, proximally located lincRNA-PCG pairs showed significantly higher expression correlations compared to random lincRNA-PCG pairs from the same chromosome (Binomial tests, p < 0.001 for all species; Fig. 4A). Positive correlations significantly outnumbered negative correlations among proximal lincRNA-PCG pairs across all species (Wilcoxon rank sum test, p ≤ 2.2e-16; Fig. 4B), indicating that lincRNAs are more likely to act as positive regulators of PCG expression in *Aspergillus*. We observed a clear distance-dependent effect, even within the 15kb *cis*-association window that we measured. LincRNAs-PCG pairs separated by <2 kb showed significantly stronger co-expression than those located 2-5 kb apart, and correlations declined progressively with increasing genomic distance (e.g. 5-10 kb, 5-10 kb; Fig. 4C). This pattern was consistent across the genus and was independent of whether the lincRNA was conserved or not (Fig. 4D).

**Figure 4.**
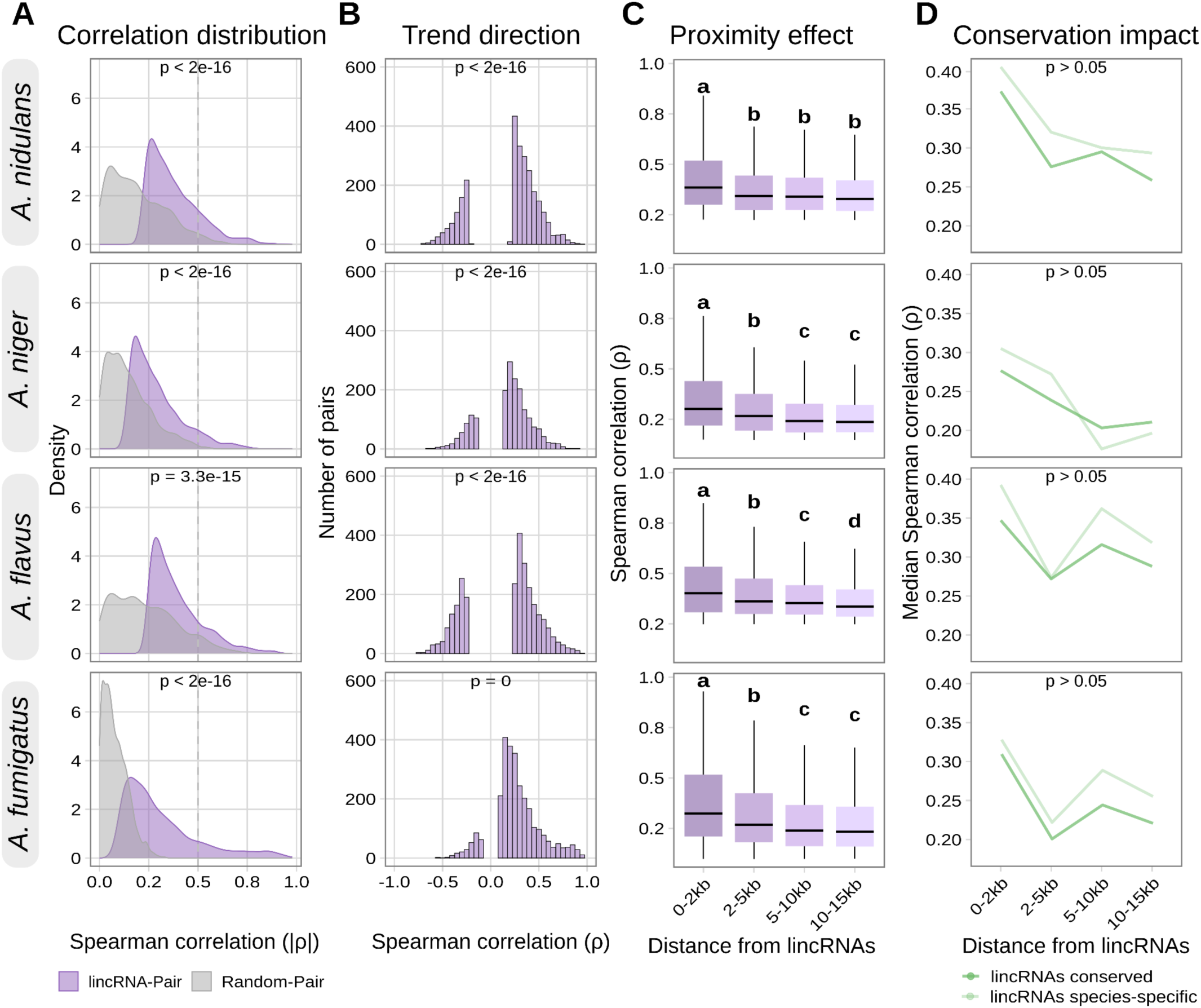
Intergenic lncRNAs display distance-dependent co-expression with nearby protein-coding genes across four *Aspergillus* species. (A) Distribution of absolute Spearman correlation coefficients (|ρ|) for lincRNA-PCG pairs within 15 kb genomic proximity and random lincRNA-PCG pairs from the same chromosome. Statistical significance assessed via binomial test. (B) Histogram showing the directionality (positive vs. negative) and correlation strength between lincRNA-PCGs pairs where p<0.05. (C) Distance-dependent decay of the absolute correlation strength (|ρ|) between lincRNAs and nearby genes. Statistical significance measured by Wilcoxon rank-sum with Benjamini-Hochberg correction. Letter groupings indicate statistically distinct groups at p < 0.05, where groups sharing a letter are not significantly different. (D) Median correlation values stratified by genomic distance and lincRNA conservation status (species-specific vs. conserved, *i.e.* present in ≥2 species). Within each species, statistical significance calculated for each distance by Wilcoxon rank-sum test. p was greater than 0.05 for all comparisons within a species.

Although we found compelling evidence for *cis*-assocations, we also wanted to explore potential long-range or *trans*-acting roles whose regulatory influence extended across distant genomic regions or chromosomes and reconstructed global co-expression networks using WGCNA [35,43]. Species-specific networks displayed typical scale-free topologies (R² > 0.8; Suppl. Fig. S3), characterized by a small number of highly connected hub genes and many nodes with few connections, a structure that distinguishes biologically organized networks from random associations [44]. Although protein coding genes unsurprisingly showed higher connectivity, lncRNAs were consistently integrated across diverse modules in all four species, rather than being isolated to a small minority of modules or those with no clear biological function based on their membership (Suppl. Fig. S4A-C). Notably, the degree of lncRNA integration varied significantly between species. In *A. flavus*, lncRNAs were members of 95% of all identified modules (39/41), while *A. fumigatus* exhibited the most selective integration of lncRNAs into the global gene expression network, with only 74% of modules containing ≥1 lncRNA (26/35) (Suppl. Table S5). Altogether we find evidence that lncRNAs likely act largely in *cis* to regulate nearby protein coding genes and are well integrated into the global gene expression network.

### LncRNA integration reflects species-specific specialization

Given that most co-expression modules contained at least one lncRNA, we next wanted to use their widespread participation in the gene expression network to predict putative biological roles for the lncRNAs described in our study. For this, we applied a guilt-by-association framework, where strong co-expression with protein coding genes of known function was used to assign putative lncRNA functions. Of the 8,553 lncRNAs discovered across the four species, 30% were assigned to co-expression modules (n=2,819) (Suppl. Table S5). Using GO enrichment of co-expressed PCGs, we inferred putative biological roles for 2,291 lncRNAs (81% of module-assigned transcripts; Suppl. Fig. S4). The unique lncRNA-enriched modules for each species reflected their divergent specializations (Suppl. Table S5). In the opportunistic pathogen *A. fumigatus*, lncRNAs were concentrated in modules governing nutrient scavenging, metal homeostasis, and oxidative stress; functions essential for metabolic flexibility in diverse environments such as compost and the human lung (*e.g.* bisque4 module, 28% lncRNA membership) [45–48]. In the industrial cell factory *A. niger,* lncRNAs-enriched modules were linked to complex carbon metabolism, thermal adaptation, and protein degradation (*e.g.* darkgreen module, 9% membership), processes central to biomass decomposition and fermentation. Notably, in *A. flavus,* a lincRNA emerged as a high-connectivity hub within the aflatoxin biosynthesis module. Together, these co-expression patterns provide a hypothesis-driven roadmap for functionally investigating the myriad ways in which lncRNAs regulate specific and broader metabolic and developmental programs.

### Loss of the lincRNA *aflalinc* significantly reduces aflatoxin biosynthesis

Fungi, including *Aspergilus*, synthesize a dazzling array of secondary metabolites, which are used for microbial communication and defense, but often act as toxins or virulence-associated factors during interaction with non-microbial partners. Our co-expression networks revealed that several secondary metabolism modules in multiple species contained lncRNAs members, including the salmon1 module in *A. flavus.* This module contained the aflatoxin biosynthetic gene cluster (BGC) and two newly described lncRNAs. We specifically focused on the lncRNA with the highest co-expression weight and named this transcript *aflalinc* (MSTRG.8604.5). This intergenic *aflalinc* was found only in *A. flavus* and was composed of six exons and spanned 3,447 bp in the genome. Confirming its classification as a non-coding transcript, it did not contain any detectable Pfam domains and lacked coding potential, as measured by a CPC2 probability 0.039 and FEELnc score of 0 (Suppl. Fig. S6A). Consistent with the TE-associated origin of many lincRNAs, *aflalinc* contained a 79 bp exon fragment belonging to the unclassified category of TEs. Despite being located on the opposite chromosome end and ∼4 Mb away from the aflatoxin BGC, *aflalinc* was strongly co-expressed with 18 cluster genes (Pearson *r* > 0.8), indicating a potential *trans*-acting modulation of aflatoxin biosynthesis. This positive co-expression included the backbone enzymes *aflC, aflA, aflB* and the pathway-specific regulator genes *aflR* and *aflS* (Fig. 5B, C, Suppl. Fig. S6B).

**Figure 5.**
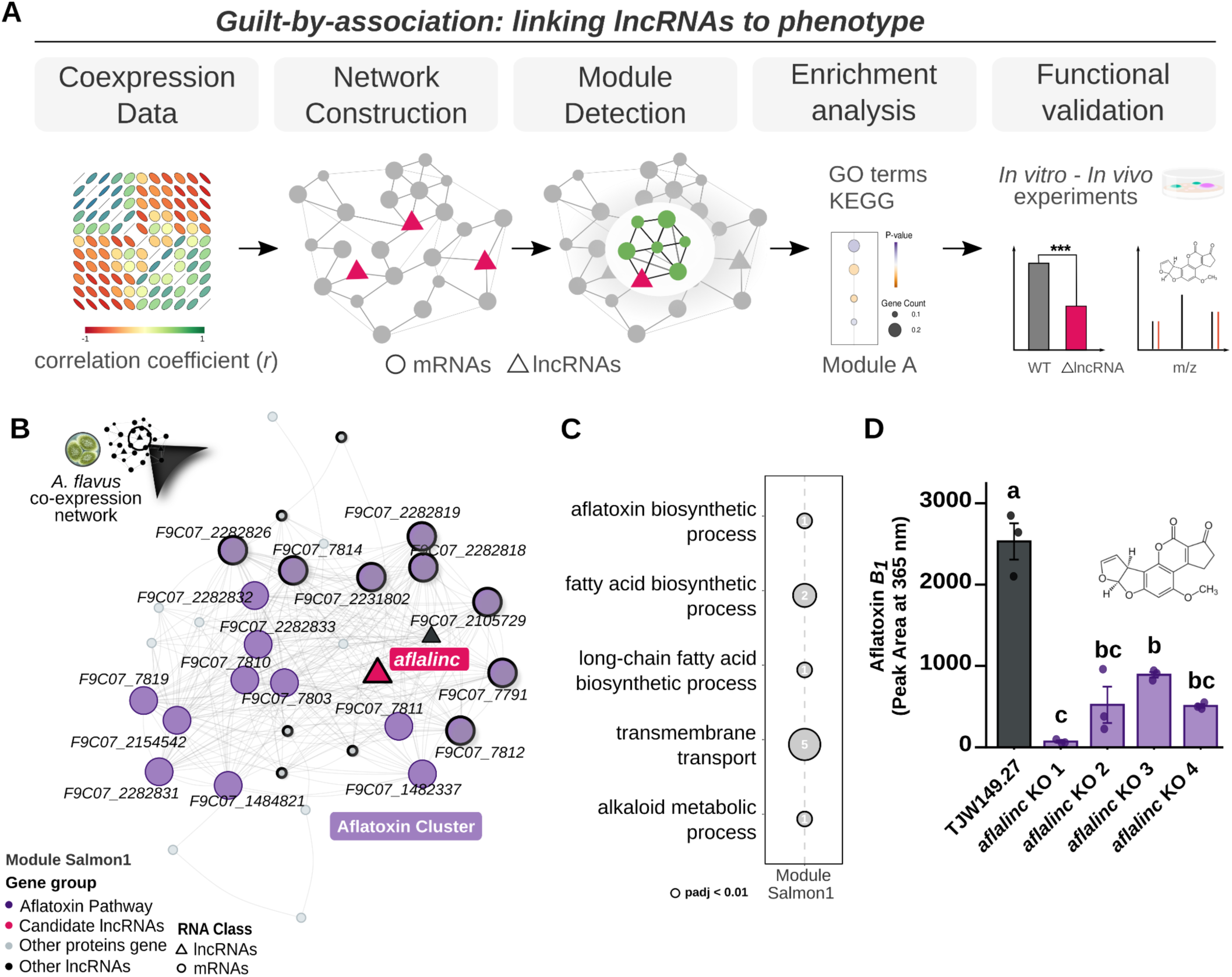
Co-expression analysis identifies *aflalinc* as a lincRNA important for aflatoxin *B_1_* biosynthesis in *A. flavus*. (A) Schematic illustrating the co-expression workflow and guilt-by-association strategy for identifying GO and KEGG functional annotations associated with lncRNAs. (B) Sub-network of the salmon1 module containing the aflatoxin BGC. Nodes represent individual genes (lncRNAs and PCGs), while edges represent high topological overlap (TOM). Genes directly connected to *aflalinc* are highlighted with black circles. *aflalinc* is identified as a high-degree hub within this module, showing strong connectivity to aflatoxin BGC genes (violet). (C) Gene Ontology (GO) terms that were overrepresented in the salmon1 module. Statistical enrichment was determined by Fisher’s exact test with Benjamini-Hochberg correction (p < 0.05). Numbers indicate the number of protein coding genes in the Salmon1 module that carry a given GO annotation. (D) Experimental quantification of aflatoxin *B_1_* production in the TJW149.27 parent strain and four independent Δ*aflalinc* mutants. Aflatoxin *B_1_* levels were measured by HPLC and the bar heights represent the mean of three biological replicates, with error bars indicating the standard error of the mean (SEM). Individual data points (n=3) are displayed for each strain.

Given the public-health importance of aflatoxin *B₁* as a potent agent of acute toxicity and global driver of hepatocarcinoma, we asked whether the co-expression between the BGC and *aflalinc* was functionally significant. For this, we generated four independent deletion mutants of *aflalinc* and confirmed them by Southern blot (Suppl. Fig. S6C). Remarkably, all four *Δaflalinc* mutants had significantly reduced aflatoxin *B₁* production levels when levels were measured by HPLC (Fig. 5D). Importantly, the phenotypic effects observed were also specific to aflatoxin biosynthesis. The *Δaflalinc* mutants did not show any phenotypic alterations in colony morphology, growth, conidiation, or sclerotia biomass formation (Suppl. Fig. S6D-F), indicating that *aflalinc* is dispensable for growth under basal conditions and instead contributes specifically to modulating aflatoxin biosynthesis. Together, our comparative analysis of lncRNAs in *Aspergillus* identified *aflalinc* as a novel, species-specific regulator of aflatoxin biosynthesis, pointing to it as a promising target for interventions reducing aflatoxin contamination of food crops.

### Analysis of an independent dataset validates co-expression network robustness and reveals stage-specific roles for lincRNAs during *A. fumigatus* biofilm maturation

Finally, we wanted to investigate the potential contributions of non-coding genes in the context of human pathogenicity and infection. For this, we turned to a previously generated transcriptome dataset of *A. fumigatus* biofilm maturation, a central virulence trait in invasive infection [49,50]. Because our initial *A. fumigatus* co-expression network was not built using this dataset or any other biofilm data, it also allowed us to assess the robustness of the gene co-expression modules against a condition not seen during network construction. When we projected the differentially expressed genes from the independent *A. fumigatus* biofilm maturation dataset onto the *A. fumigatus* global co-expression network, we observed that 25 of 35 modules showed moderate to high preservation (Zsummary > 2; Suppl. Fig. S5, Suppl. Table S6) (Fig. 6A). This demonstrates that the identified co-expression relationships persist regardless of whether the condition was present during network construction.

**Figure 6.**
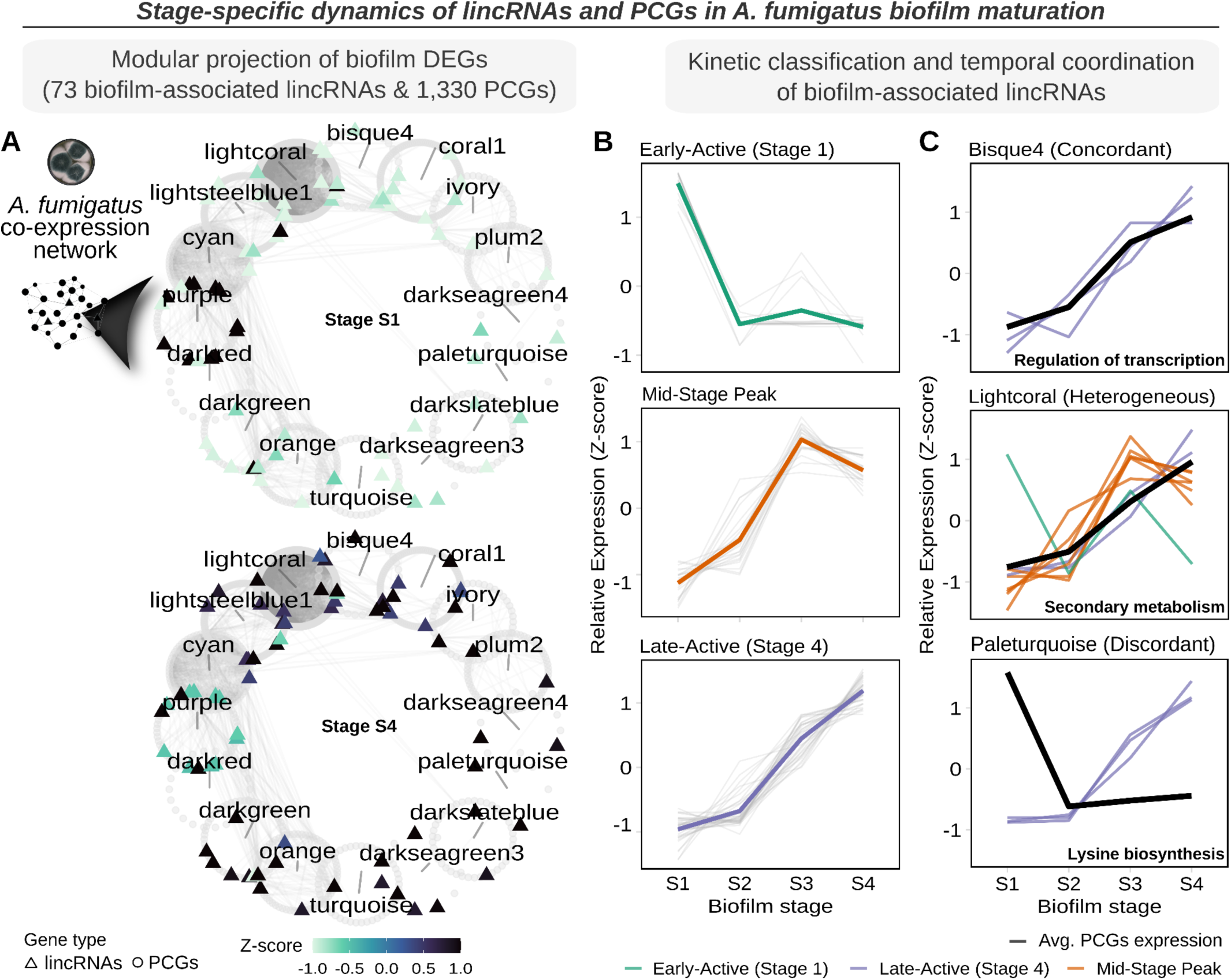
Dynamic expression of lincRNAs across biofilm stages in *Aspergillus fumigatus*. (A) Representation of the co-expression modules containing non-coding and protein-coding genes differentially expressed during biofilm maturation (|log2FC| > 1.5, padj < 0.05). Node shape distinguishes protein-coding genes (○) from lincRNAs (▵), while node color indicates Z-score normalized expression. Only modules containing lincRNAs are displayed. Edges represent co-expression links between lincRNAs and protein-coding genes within and across WGCNA modules. (B) Temporal dynamics of lincRNAs across biofilm stages. Individual lincRNAs trajectories are shown as faint gray lines, with bold colored lines representing the average expression of each temporal group: Early-Active (peak at S1/12 hr), Mid-Stage Peak (S2-S3 / 18-24 hr), and Late-Active (S4/30 hr peak). (C) lincRNAs-PCGs temporal relationships within individual modules. Black lines show average protein expression for the module; colored lines indicate individual lincRNAs trajectories colored by temporal classes (teal: early, orange: mid, lavender: late).

The dataset is composed of four timepoints, representing early (12 and 18 hour) and late (24 and 30 hour) stages of biofilm architecture and maturation. To define the novel participation of lncRNAs relevant to pathogenesis, we focused on the 73 intergenic lincRNAs that were differentially expressed in at least one time point (|logFC| > 1.5, FDR adjusted p < 0.05) and were assigned to 16 co-expression modules. Based on expression trajectory over the time course, transcripts were partitioned into three groups: Early-Active (peaking at Stage 1/12 hr), Mid-Stage Peak (Stage 2-Stage 3; 18 and 24 hr), and Late-Active (peak at Stage 4/30 hr) (Fig. 6B, Suppl. Table S6). While 11 modules showed synchronous timing, we identified five discordant modules where lincRNA activity preceded or opposed protein expression (Fig. 6C). As lncRNA levels often peak before their protein partners as part of their regulatory roles, these lincRNAs may act as early signals that prepare the module for subsequent stages of development. In the secondary metabolism module (lightcoral), which includes pathways such as pseurotin A, fumagillin, and fumitremorgin B, lincRNAs peaked during mid-stages (S2–S3) while proteins peaked late (S4). Conversely, in the paleturquoise module (lysine biosynthesis), lincRNAs peaked late (S4) while proteins peaked early (S1). Even within concordant modules like purple (iron homeostasis), lincRNAs overwhelmingly peaked early (79% S1) compared to the broader trajectory distribution of protein-encoding transcripts. Altogether, this temporal catalog of 73 biofilm-associated lincRNAs demonstrates the robustness of the co-expression networks and serves as a starting point for probing how non-coding transcripts coordinate biofilm and other pathogenicity programs.

## Discussion

In this study we establish a comprehensive catalog of 8,553 lncRNAs spanning a large swath of the genus *Aspergillus*, providing insight into the non-coding RNA “dark matter” in fungal genomes. Annotation of lncRNAs expanded the gene content of each species by up to 20% and provides a platform for systematically interrogating the roles of non-coding genes in this medically and economically important genus. Of this lncRNA catalogue, only 488 intergenic gene families were conserved across more than one species, highlighting lncRNAs as a source of genomic novelty and plasticity across *Aspergillus* spp. Notably however, the lincRNAs that were conserved were expressed much more strongly and were implicated in central functions based on co-expression. We find evidence that the abundant magnitude of lncRNAs is driven at least in part by transposable elements, as TE sequence is overrepresented in lncRNAs–a finding particularly true for subtelomeric lncRNAs. Through combined analysis of *cis* and global co-expression, we find that lncRNAs most commonly show a stronger association with nearby genes, but our comparative analyses also identified *aflalinc* as a novel lincRNA in *A. flavus* that acts in *trans* to modulate aflatoxin production. When deleted, aflatoxin production was significantly reduced, opening the door for targeted, species-specific interventions. Together this work provides the first experimental demonstration that a lncRNA can strongly and specifically modulate BGC activity in *Aspergillus* and establishes that our *in silico* approaches capture genuine functional associations.

Our work reveals that lncRNAs in *Aspergillus* are strongly intertwined with transposable elements, with up to 45% of lincRNAs in *A. fumigatus* containing TE-derived sequences. As ours is the first study to comprehensively calculate the link between TEs and lncRNAs in the fungal kingdom we have no point of comparison, but we expect follow-up studies will reveal connections in other fungal species, based on observations in other eukaryotes [10–12]. Unlike PCGs, where TE insertions are purged by purifying selection, lincRNAs in *Aspergillus* appear to readily accommodate these elements. This is consistent with the RIDL hypothesis [13], where TE fragments are leveraged as pre-built functional blocks for nascent lncRNA functions. The absence of transposase signatures and the high proportion of unclassified mobile genetic elements in *Aspergillus* lncRNAs points to decayed TEs; something that could be aided by repeat-induced point mutation (RIP) [12,40,51–54]. While the intrinsic goal of RIP is to mutagenize and silence multicopy repetitive loci such as TEs, in the context of lncRNAs it might function as a structural architect, converting mobile elements into stable non-coding exons. These TE-associated lincRNAs are preferentially located in subtelomeres, regions known in fungi as safe harbors for adaptive protein-coding genes and BGCs [55,56], and are expressed at very low levels, suggesting additional epigenetic containment [41]. We propose that, in *Aspergillus*, subtelomeres serve a dual role: they not only harbor adaptive PCGs but also act as nurseries for lncRNA innovation, where TE-driven structural expansion is tolerated, but restrained under heterochromatic control.

Synteny-based conservation identified only 488 conserved lincRNA families, indicating a huge amount of variation in the non-coding repertoire of *Aspergillus*. However, these conserved transcripts were expressed at significantly higher levels than species-specific ones. This expression hierarchy suggests functional importance, as these lncRNAs presumably originated 62-77 million years ago depending on the species pairing and have been retained throughout evolutionary time despite rapid sequence evolution among lncRNAs. Although synteny-based approaches are widely used to infer lncRNA conservation, they are inherently sequence-agnostic and can both underestimate conservation of rapidly evolving lncRNAs and overestimate conservation when non-homologous transcripts arise within conserved genomic intervals [11,57]. Regardless of these methodological considerations, the stronger co-expression between lincRNAs and nearby PCGs points to local *cis*-associations, concordant with patterns reported in mammals [58–60]. Although co-expression does not establish causality, it is consistent with a role in local transcriptional regulation that may contribute to the evolutionary retention of the lncRNAs. Even though PCGs are much more stable than lncRNAs, it is becoming increasingly appreciated that there is still considerable variation in presence or absence of PCGs in isolates even within the same species. For *A. flavus* and *A. fumigatus* where this has been systematically quantified using large datasets, only 57-69% of genes are conserved species-wide [42,61]. Given the rapid evolution of lncRNAs, we expect that the proportion of non-coding genes conserved between isolates to be markedly lower. The exact degree of lncRNA variation within the same species and their roles in isolate-specific phenotypes will be an interesting area to explore in the future pending the availability of transcriptomes from a large and diverse strain panel.

The identification of *aflalinc* as a strong modulator of aflatoxin biosynthesis and whose deletion produces one of the most pronounced non-coding phenotypes reported in filamentous fungi to date opens up many exciting avenues for future work. Unlike the cluster-resident regulators *aflR* and *aflS*, *aflalinc* operates from a distance of ∼4 Mb, underscoring the ability of co-expression networks to reveal long-range associations invisible to proximity-based searches. We also expect that the modulation of aflatoxin by *aflalinc* is independent of the global secondary metabolite regulator *laeA* [62], as no connections were observed between the two in the co-expression network. Further, the deletion of *aflalinc* led to marked reductions in *AFB₁* production without affecting growth or conidiation, whereas *laeA* deletion impacts all three [63]. Moreover, it was not found in the other *Aspergillus* spp examined, indicating it could be targeted specifically without affecting other aspects of *A. flavus* growth, other non-pathogenic *Aspergillus* spp. present or the endogenous mycobiome. These together make *aflalinc* a promising target for spray-induced RNAi silencing, a field that has shown great promise against other plant pathogens [64–66] with initial results supporting this method for aflatoxin control in transgenic maize [67,68].

While the specific mechanism by which *aflalinc* works to modulate aflatoxin expression remains unknown, several possibilities emerge. One is that *aflalinc* influences chromatin accessibility at the aflatoxin locus, potentially by recruiting or modulating epigenetic complexes such as COMPASS (Bre2) or the chromatin remodeler Arp9, both recently implicated in aflatoxin regulation [69,70]. Alternatively, *aflalinc* may act as a decoy, sequestering repressor proteins that block the one or more factors important for aflatoxin production [27]. Both possibilities would be consistent with the major mechanistic modes of lncRNAs, which are guiding chromatin modifiers, scaffolding regulatory complexes, or titrating away inhibitory factors and would all be compatible with its long-range influence on the aflatoxin BGC. Defining these interactions will require ChIRP-MS to identify its protein partners and ChIRP-seq or CHART-PCR to map its direct binding sites on DNA, allowing the determination of whether *aflalinc* physically associates with promoters or regulatory elements within the aflatoxin cluster.

Altogether, our work establishes a comprehensive resource for future work identifying the roles by which non-coding transcripts contribute to fungal development and metabolism. Aflatoxin biosynthesis in *A. flavus* and biofilm maturation in *A. fumigatus* represent two fungal traits with major consequences for human health, one driving global food-safety crises, the other underlying invasive disease in immunocompromised patients. By examining lncRNAs across these distinct contexts, we show that non-coding transcripts participate in both mycotoxin-associated metabolism and are linked to biofilm development programs. Despite the strengths of our approaches, most predicted lncRNAs still require experimental validation, representing a major experimental bottleneck. However, high throughput functional genomic advances such as multiplexed CRISPR interference and activation can complement the *in silico* analyses used here to further prioritize lncRNA candidates for individual functional validation [71]. Ultimately, this study represents a major step toward defining the unique non-coding transcriptomes of the genus *Aspergillus* and provides myriad opportunities for interventions and future work relevant to food safety, biotechnology, and human pathogenicity.

## Materials and methods

### Reference Genomes and Annotation

Reference genomes (FASTA format) and corresponding annotation files (GFF3 format) for the four *Aspergillus* species, *A. fumigatus* (Af293)*, A. flavus* (NRRL3357)*, A. niger* (CBS 513.88), and *A. nidulans* (FGSC A4), were downloaded from the 68 release of FungiDB [72,73]. We performed the lncRNA prediction independently for each of the four species. To supplement the publicly available annotations and standardize the prediction of biosynthetic gene clusters (BGCs), secondary metabolites were re-annotated using the fungal version of antiSMASH v7.1.0 [74]. Gene Ontology (GO) terms and functional annotation were assigned using the Gene Annotation Format (GAF) files from FungiDB (release 68, accessed in November 2024). Protein domain annotations (PFAM) and functional predictions were obtained using InterProScan v5.68-100.0 [75].

### Public RNA-seq data collection

We compiled RNA sequencing data for *A. fumigatus, A. nidulans, A. niger,* and *A.flavus* from the NCBI Sequence Read Archive (SRA) database (Suppl. Table S1). We retrieved and selected only those BioProjects with paired-end and stranded sequencing using sra-tools v3.0.10 (https://github.com/ncbi/sra-tools) with prefetch and fastq-dump functions. To infer strand specificity for each sample without prior metadata we applied the same methods as in [35]. Briefly, we used Salmon v0.8.1 in pseudo-mapping mode [76]. Quantification was performed on trimmed paired-end reads against a pre-built transcriptome index derived from the appropriate *Aspergillus* reference genome for each species. Salmon was executed with the -l A parameter, which enables automatic inference of library type by examining k-mer orientation and fragment mapping direction. The resulting strand information was used to guide accurate transcript assembly in downstream steps.

### RNA-seq mapping and transcriptome assembly

For reproducible and scalable pre-processing, we implemented a custom workflow using Snakemake v7.30.1 [77], a python-based workflow management system. For quality control of the raw FASTQ files, FastQC v0.12.1 (https://github.com/s-andrews/fastqc) was used and results aggregated with MultiQC v1.15 [78]. Next, we continued with adapter and quality trimming: reads were trimmed using fastp v0.23.4 with customized parameters to ensure high-quality read retention [79]. Specifically, we applied 5′ and 3′ end trimming (-cut_front 3--cut_tail 3), removed poly-G and poly-X tails (--trim_poly_g --trim_poly_x), enforced a minimum read length of 49 bp (--length_required 49), and filtered reads based on sliding window quality (-W 4 -M 5). Trimming was performed in paired-end mode with multithreading support. Reads passing QC were aligned to the respective *Aspergillus* reference genomes using STAR v2.7.10b with splice-aware mapping [80]. The alignments were sorted by genomic coordinates (--outSAMtype BAM SortedByCoordinate), and splice junction annotations were provided via GTF files (--sjdbGTFfile). To accommodate fungal genome structure, the maximum intron length was limited to 2000 bp (--alignIntronMax 2000). Finally, transcriptome assembly was performed using StringTie v2.1.7 in reference-guided mode (-G), where reads were assembled into transcript models [81]. To enhance the specificity of transcript prediction, we applied a minimum transcript coverage threshold of 1.5 (-c 1.5) and a minimum isoform abundance fraction of 0.05 (-f 0.05). Strand-specific assembly was performed based on library type, using the strandness information derived from the Salmon output.

### *In silico* lncRNA identification

For lncRNA prediction, we followed the methods outlined by Hovhannisyan *et al.* (2022). In brief, we merged all the assembly samples to create a unified, non-redundant transcriptome for each *Aspergilli* species, using the Stringtie merge function with the -g 50 option. We then compared the transcriptome annotations with the original genome annotations using gffcompare v0.12.6 [35,82]. This allowed us to identify novel transcripts classified as intergenic lncRNAs (lincRNAs), which are in the genomic interval between genes, and antisense lncRNAs (As lncRNAs), which overlap with protein exons but are transcribed from the opposite strand. We used CGAT gtf2gtf v0.6.0 software to ensure that only the longest isoform was retained when multiple options were available [83]. Following that, we assessed the coding potential of these predicted transcripts with CPC2 v1.0.1 and Feelnc v0.2 (shuffle mode) software and kept only the transcripts classified as “non-coding” by both programs [84,85]. Finally, we performed multiple filtering steps to generate our final putative lncRNAs catalog for downstream analysis. First, we used cgatcore v0.6.15 to keep only those non-coding transcripts longer than 200 bp and >1 TPM in at least 2 samples [83]. Second, and only for the intergenic lncRNAs, we assessed homology to known protein domains using BLASTX (BLAST+ v2.9.0) with parameters set to e-value < 1e-5, minimum query coverage ≥ 50%, and percent identity > 30, against the SwissProt database (downloaded July 2024) [86]. Finally, structural non-coding RNAs such as tRNAs, rRNAs, and snoRNAs were identified using Infernal v1.1.4 with the cmscan program against the Rfam covariance model (Rfam.cm, with clan file support and --cut_ga thresholds) [87,88]. Transcripts matching Rfam entries or showing protein domain hits were flagged for exclusion. To integrate novel lncRNA annotations into the existing reference genome annotations of four *Aspergillus* species (*A. flavus, A. fumigatus, A. nidulans,* and *A. niger*), we first converted the GTF file containing the filtered lncRNA transcripts (retaining only the longest isoform per gene and preserving exon structure) into GFF3 format using GffRead v0.12.7 [82]. Subsequently, we merged the resulting lncRNA GFF3 annotation with the corresponding reference genome annotation of each species using AGAT v0.8.1 (agat_sp_merge_annotations.pl)[89].

### Intergenic lncRNAs conservation analysis

To assess the evolutionary conservation of intergenic lncRNAs across *Aspergillus* species, we created a synteny-based strategy based on the methodology outlined in Pegueroles et al. (2019 [38]). First, we identified one-to-one orthologous genes across the four species using the GENESPACE R package v1.3.1 [90]. This tool clusters protein sequences into orthologous groups using OrthoFinder v2.5.5 [91], followed by a synteny-aware refinement to resolve multicopy ortholog groups where possible. We then extracted lincRNAs from each species based on sequence IDs (containing the tag |u|) in species-specific BED files using BEDTools (v2.31.1) [92]. For each lincRNA, the pipeline retrieved three flanking genes on each side, defining a syntenic neighborhood. Syntenic conservation between species was defined as a minimum overlap of flanking genes on each side of the lincRNA and one additional shared gene in this neighborhood, (parameters: genesNearby = 3, minOverlap = 3, minSideOverlap = 1). Pairwise comparisons among the four species were conducted in both directions to capture reciprocal conservation. The output of the synteny comparisons was then parsed to group homologous lincRNAs into conserved families (Suppl. Table S3).

### Comparative expression and chromosomal distribution of protein-coding and non-coding genes

To assess gene expression variability across different gene types, we obtained the raw read counts from RNA-seq alignments (BAM files) using featureCounts v2.0.1 [93] and determined the Transcripts Per Million (TPM) values. These normalized expression values were used to compute the coefficient of variation (CV) for assessing gene expression variability across different gene types. To investigate the genomic distribution of transcript types (PCGs, lincRNAs, and As lncRNAs), we generated chromosome-level binned count plots per species. Chromosome sizes were extracted from the reference genome annotation file (GFF3). We adopted the method of Zhang et al. (2024 [41]) to define subtelomeric regions as the terminal 10% of each chromosome. These coordinates were identified using custom R scripts and BEDTools (v2.31.1) [92]. We then assessed overlap between transcript coordinates of lncRNAs and protein-coding genes with the defined subtelomeric regions through the intersectBed function. A transcript was classified as subtelomeric if any part of its genomic span intersected with our defined subtelomeric region. To evaluate the enrichment of lncRNAs or protein-coding genes, we constructed a 2 × 2 contingency table for each transcript category. We employed a Chi-square test of independence to determine whether the distribution of transcripts differed significantly from a random distribution across the chromosomes.

### Identification of transposable elements and their overlap with lincRNAs

Transposable elements were annotated in each *Aspergillus* spp. genome using EarlGrey (v4.3.0; *earlGreyAnnotationOnly*())[94]. The non-redundant transposable element database mycoMobilome (v1.0) was used as the custom TE library for RepeatMasker [51]. Specifically, the consensus sequences supported by protein evidence (_PE) were utilized to ensure high-confidence mapping of ancestral and potentially active TE families across all four species. The resulting TE annotations were categorized into major TE families, including DNA transposons, long terminal repeat (LTR) retrotransposons, long interspersed nuclear elements (LINEs), short interspersed nuclear elements (SINEs), unclassified, and other repeat elements. To identify lncRNAs overlapping TEs, we used the intersectBed function from BEDTools (v2.31.1).

Intergenic lncRNA were classified as TE-associated if at least one exon contained ≥10 bp TE sequence. In cases where multiple TE fragments overlapped the same transcript, the lincRNA was considered a single TE-associated unit. The TE fraction was calculated as the total base pairs of all contained TE sequences divided by the total transcript length. Queries to assess whether any lincRNAs were potentially misannotated, but functional TEs was performed using BLASTX (BLAST+ v2.9.0) against a curated database of transposon-related proteins (Transposase, Reverse Transcriptase) and using relaxed search criteria of a maximum E-value of 1e-5, coverage > 50%, and identity > 30%. No BLASTX hits fulfilling these criteria were returned, confirming the absence of functional enzymes for transposition or reverse transcription among lincRNAs.

### Correlation of *cis*-lincRNAs genes

Coordinates of lincRNAs and protein-coding genes were extracted from BED files for each *Aspergillus* species. Genomic ranges were constructed using the GenomicRanges R package v1.58.0 [95] to identify protein-coding genes located near intergenic lncRNAs, which were defined as genes within a 15 kb window of either the start or the stop of the lincRNA, irrespective of transcriptional direction. Strand orientation and distances were calculated, and overlaps were identified using the *findOverlaps* function. To evaluate co-expression, Spearman Rank’s correlation coefficients were calculated in R between the normalized expression values of each lincRNA and its nearby genes across all samples. Correlation distributions were compared across different genomic distance bins (e.g., ±1 kb, ±5 kb, ±10 kb), as well as the same number of with randomly paired lincRNA-PCGs pairs located on the same chromosome, which served as a background model. Statistical significance between correlation distributions was assessed using the Binomial test and Wilcoxon rank-sum test.

### Co-expression network analysis

To identify co-expression modules involving lncRNAs and PCGs, we performed a weighted gene co-expression network analysis (WGCNA) using the R package WGCNA v1.73 [43]. This analysis was performed separately for each *Aspergillus* species. The raw counts expression matrices were normalized to TPM and then log2(TPM +1) was used as the input. To reduce background noise and focus on more consistently expressed genes, only those with an expression level above 0.1 TPM in at least 80% of the samples were retained. Before constructing the network, we determined the appropriate soft-thresholding power (β) for each species using the *pickSoftThreshold* function from the same package. The objective was to identify the smallest power at which the network approximates scale-free topology (R² ≥ 0.80) while maintaining sufficient connectivity [44]. The selected soft-thresholding powers were: β = 10 for *A. flavus*, β = 6 for *A. fumigatus*, β = 12 for *A. nidulans*, and β = 7 for *A. niger*. These values were applied in all subsequent steps to construct the unsigned adjacency using the *wgcna::cor*() function. Next, the adjacency matrix was transmutated into topological overlap measure (TOM) values, which was then used to perform hierarchical clustering based on the TOM dissimilarity measure (1-TOM). Modules of co-expressed genes were identified using dynamic tree cutting (*cutreeDynamic*) with deepSplit set to 3 and a minimum cluster size of 20 genes. We calculated module eigengenes and merged similar modules at a correlation threshold of 0.75 (height = 0.25). We then assigned each gene to a final co-expression module. In the WGCNA framework, each module is automatically assigned a unique color label for identification. Intramodular connectivity (kWithin) was computed to evaluate the centrality of genes within their respective modules using the *intramodularConnectivity* function within the R package WGCNA. Network visualizations were done with tidygraph v1.3.1 and ggraph v2.2.1 [96,97].

### Inferring lncRNA function through guilt-by-association

To characterize the biological properties of the identified co-expression modules, we performed Gene Ontology (GO) enrichment analyses on the PCGs within each module using clusterProfiler v4.14.6 [98]. Enrichment was conducted separately for the Molecular Function (MF), Biological Process (BP), and Cellular Component (CC) ontologies of GO using the enricher function with species-specific gene sets as the background. Significantly enriched terms (FDR-adjusted p < 0.05) were identified for each of the MF, BP, and CC domains. The results from GO term analyses, along with the list of associated lncRNAs, were compiled into a unified annotation table for each module to facilitate cross-species comparisons (Suppl. Table S5). Modules were manually assigned to broad functional categories based on their enriched GO terms and literature review. Categories included “Pathogenesis & Nutrient Scavenging”, “Specialized Toxins & SM”, “Growth & Development”, “Industrial & Carbon Metabolism”, and “Stress & Resilience” (Supplementary Table S5). For each category, we calculated the “regulatory investment” as the total number of lncRNAs in modules assigned to that category divided by the total number of genes in those modules. This normalizes for module size and reflects the proportion of non-coding transcripts within a given functional domain.

### Biofilm transcriptome analysis

For identification of lncRNAs involved in *A. fumigatus* biofilm establishment and maturation we examined the RNA-seq datasets generated in [50] and deposited into the NCBI SRA under PRJNA1269323. Quality control and mapping of transcriptome data were performed as described above. Genes with fewer than 10 total counts across all samples were filtered out prior to analysis. Differential expression analysis was performed using DESeq2 v1.46.0 [99]. Pairwise comparisons were conducted between Stage 1 (initial colonization, 12 h) and Stage 4 (mature biofilm, 36 h). Genes were considered differentially expressed (DEGs) if their false discovery rate (FDR) was < 0.05 and absolute log2 fold change > 1.5. To map biofilm dynamics onto our previously constructed co-expression framework, we projected the DEGs, proteins and lincRNAs, onto the global *A. fumigatus* WGCNA network. The TOM was thresholded at 0.10 to remove weak edges and enhance visualization clarity. To assess temporal dynamics, expression values for each gene were Z-score normalized across all 12 samples, centering each gene’s trajectory at zero to enable cross gene comparisons. To categorize lincRNA expression patterns, we developed a R custom script with a classification function that assigns each lincRNA to one of three categories based on its peak activity stage: Early-Active (peak at Stage 1), Mid-Stage Peak (peak at Stage 2 or 3) and Late-Active (peak at Stage 4). This classification was used to group lincRNAs with similar temporal dynamics and to assess stage-specific enrichment within modules.

### Generation of the *aflalinc* (Δ*MSTRG.8604.5*) deletion strains

To investigate the functional role of *aflalinc* (Δ*MSTRG.8604.5*), deletion mutants were generated in an *Aspergillus flavus* TJES19.1 (NRRL3357 *ΔpyrG*, *ΔKu70* [100]) background. A deletion cassette was constructed using a double-joint PCR strategy [101]. Briefly, 1 kb genomic DNA fragments flanking the 5′ and 3′ regions of the *aflalinc* locus were amplified from *A. flavus* NRRL3357 and fused to a 2 kb pyrG selectable marker amplified from *A. fumigatus* Af293. Fungal transformation was performed as previously described [62,102]. Transformants were initially screened by PCR and subsequently validated via Southern blot analysis. Genomic DNA from the parental and mutant strains was digested with *Eco*RI and hybridized with ^32^P-labeled probes corresponding to the 5′ and 3′ flanking regions to confirm the targeted replacement of the native locus and the absence of ectopic integrations. A total of four independent sibling mutants were selected for further phenotypic and metabolic analysis. Strains and primers are listed in Suppl. Table S7.

### *A. flavus* strains and growth conditions for fungal growth and secondary metabolic assays

*A. flavus* strain TJW149.27 (NRRL3357 *Δku70, pyrG+,* [103]) and the four Δ*aflalinc* deletion mutants were grown on glucose minimal medium (GMM). Spores were harvested from 5-day-old cultures and adjusted to identical concentrations. For colony morphology and metabolite production, 10^2^ spores were plated on GMM supplemented with 2% sorbitol using an overlay technique with 0.75% top agar. Cultures were incubated at 30°C in the dark for 7 days prior to analysis.

Conidiation was quantified by collecting three agar plugs (totaling 1 cm²) per plate into 3 mL of 0.01% Tween 80. Spores were dislodged through a sequence of vortexing (1 min), sonication (30 s), and a final vortexing step (30 s). Spore concentrations were determined using a Cellometer X2 with Cellometer Image Cytometer software (v3.2.1; Nexcelom Bioscience LLC). For sclerotia quantification, sclerotia were mechanically harvested from the plate surface using a metal rod. Samples were transferred to pre-weighed microcentrifuge tubes, lyophilized for 48 hours to ensure complete dehydration, and weighed to determine total sclerotial biomass per plate. All experiments were performed with four independent biological replicates.

### Metabolite extraction and aflatoxin analysis

Aflatoxin was extracted from the agar media using a modified organic solvent extraction protocol [104]. Media were pulverized and subjected to two sequential extractions with 25 mL of chloroform. Following overnight extraction at room temperature, the organic phase was filtered through Whatman No. 1 paper and evaporated to dryness under a fume hood. Aflatoxin *B_1_* was detected and quantified using analytical reversed-phase high-performance liquid chromatography (HPLC). The system consisted of a Gilson 322 pump coupled to a Gilson Verity 1741 UV–VIS detector (Gilson, Middleton, WI, USA), equipped with an Eclipse Plus C18 column (5 µm, 19 × 250 mm; Agilent, Santa Clara, CA, USA). The mobile phase utilized a gradient of 0.1% formic acid in water (A) and 0.1% formic acid in acetonitrile (B). The gradient profile was: 30% B for 30 min, linear increase to 100% B over 10 min, decrease to 30% B over 1 min, and a final hold at 30% B for 10 min (total run time: 51 min). Aflatoxin *B_1_* was detected at 365 nm with a flow rate of 1 mL/min (retention time ≈ 18.97 min). Dried extracts were resuspended in HPLC-grade methanol to a concentration of 10 mg/mL, and 20 µL was injected per sample. Quantitative comparisons were performed based on the area under the curve (AUC) relative to an aflatoxin *B_1_* analytical standard. All experiments were performed with three independent biological replicates.

### Statistical analyses

Statistical and all individualized data analyses were carried out using R v4.4.3 (R Core Team 2024) and Python v3.12.0. Specific methods and tests applied in each analysis are detailed in the relevant sections. Where multiple comparisons were involved, Benjamini-Hochberg correction was applied to control for false positives. Visualizations plots were done using ggplot2 v3.5.1 [105], ggridges v0.5.6 [106], and ggpubr v0.6.0 [107].

## Acknowledgements

Computational work in this study was performed, in part, on the HPC cluster of the Friedrich Schiller University. The authors thank Marion Perrier, Ailton Pereira da Costa Filho, and Matthew G. Blango for their scientific feedback.

## Funding

MLF was supported by a scholarship from the Friedrich Schiller University for female postdocs. The authors also acknowledge funding from the Leibniz Collaborative Excellence Program, Project K569/2023 “FuRTHER - Fungal RNA Transmission Impacting Human Epigenome Regulation” (AEB and MLF). AEB and MLF are further supported by the Deutsche Forschungsgemeinschaft (DFG, German Research Foundation under Germany’s Excellence Strategy – EXC 20151 – Project-ID 390713860). Hatch Act Formula Fund WIS05058 and United States-Israel Binational Agricultural Research Development (BARD) Fund IS-5629-23 were awarded to NPK for salary support for BJH and R01 NIH/NCCIH R01 AT 00914320 awarded to NPK for salary support of JWB. The funders had no role in study design, data collection and analysis, decision to publish, or preparation of the manuscript.

## Author Contributions

MLF and AEB conceptualized and designed the study. MLF performed formal analysis and was responsible for visualization. MLF and AEB wrote the primary draft. JWB, BJH and NPK performed experimental investigations and were involved in the editing and review of the manuscript.

## Notes

### Competing Interest Statement

The authors have declared no competing interest.

## References

1. Rutenberg-Schoenberg M, Sexton AN, Simon MD. The Properties of Long Noncoding RNAs That Regulate Chromatin. Annu Rev Genomics Hum Genet. 2016;17: 69–94. doi:10.1146/annurev-genom-090314-024939

2. Mattick JS, Amaral PP, Carninci P, Carpenter S, Chang HY, Chen L-L, et al. Long non-coding RNAs: definitions, functions, challenges and recommendations. Nat Rev Mol Cell Biol. 2023;24: 430–447. doi:10.1038/s41580-022-00566-8

3. Mudge JM, Carbonell-Sala S, Diekhans M, Martinez JG, Hunt T, Jungreis I, et al. GENCODE 2025: reference gene annotation for human and mouse. Nucleic Acids Res. 2025;53: D966–D975. doi:10.1093/nar/gkae1078

4. Avalos J, Perera-Bonaño A, Limón MC. Identification and Functions of lncRNAs in Fungi. Non-Coding RNA. 2025;11: 72. doi:10.3390/ncrna11050072

5. Gao J, Chow EWL, Wang H, Xu X, Cai C, Song Y, et al. LncRNA DINOR is a virulence factor and global regulator of stress responses in Candida auris. Nat Microbiol. 2021;6: 842–851. doi:10.1038/s41564-021-00915-x

6. Chacko N, Zhao Y, Yang E, Wang L, Cai JJ, Lin X. The lncRNA RZE1 Controls Cryptococcal Morphological Transition. Butler G, editor. PLOS Genet. 2015;11: e1005692. doi:10.1371/journal.pgen.1005692

7. Huang P, Yu X, Liu H, Ding M, Wang Z, Xu J-R, et al. Regulation of TRI5 expression and deoxynivalenol biosynthesis by a long non-coding RNA in Fusarium graminearum. Nat Commun. 2024;15: 1216. doi:10.1038/s41467-024-45502-w

8. Weaver D, Qi T, Chown H, Fraczek M, Lebedinec R, Dineen L, et al. Genome-wide discovery and phenotyping of non-coding transcripts in A. fumigatus reveals lncRNAs with a role in antifungal drug sensitivity. Nat Commun. 2026;17: 1832. doi:10.1038/s41467-026-68543-9

9. Devkota R, Poudyal N, Reyes Servin J, Lenz J, Shepardson KM, Dhingra S. A long non-coding RNA, *afu-254* , is required for the oxidative stress response, cell wall stress response, azole susceptibility and virulence in *Aspergillus fumigatus*. Microbiology; 2026. doi:10.64898/2026.01.07.698160

10. Kapusta A, Kronenberg Z, Lynch VJ, Zhuo X, Ramsay L, Bourque G, et al. Transposable Elements Are Major Contributors to the Origin, Diversification, and Regulation of Vertebrate Long Noncoding RNAs. Hoekstra HE, editor. PLoS Genet. 2013;9: e1003470. doi:10.1371/journal.pgen.1003470

11. Hezroni H, Koppstein D, Schwartz MG, Avrutin A, Bartel DP, Ulitsky I. Principles of Long Noncoding RNA Evolution Derived from Direct Comparison of Transcriptomes in 17 Species. Cell Rep. 2015;11: 1110–1122. doi:10.1016/j.celrep.2015.04.023

12. Fort V, Khelifi G, Hussein SMI. Long non-coding RNAs and transposable elements: A functional relationship. Biochim Biophys Acta BBA - Mol Cell Res. 2021;1868: 118837. doi:10.1016/j.bbamcr.2020.118837

13. Johnson R, Guigó R. The RIDL hypothesis: transposable elements as functional domains of long noncoding RNAs. RNA. 2014;20: 959–976. doi:10.1261/rna.044560.114

14. Kapusta A, Feschotte C. Volatile evolution of long noncoding RNA repertoires: mechanisms and biological implications. Trends Genet. 2014;30: 439–452. doi:10.1016/j.tig.2014.08.004

15. Lv Y, Hu F, Zhou Y, Wu F, Gaut BS. Maize transposable elements contribute to long non-coding RNAs that are regulatory hubs for abiotic stress response. BMC Genomics. 2019;20: 864. doi:10.1186/s12864-019-6245-5

16. Holdt LM, Hoffmann S, Sass K, Langenberger D, Scholz M, Krohn K, et al. Alu Elements in ANRIL Non-Coding RNA at Chromosome 9p21 Modulate Atherogenic Cell Functions through Trans-Regulation of Gene Networks. McCarthy MI, editor. PLoS Genet. 2013;9: e1003588. doi:10.1371/journal.pgen.1003588

17. Carlevaro-Fita J, Polidori T, Das M, Navarro C, Zoller TI, Johnson R. Ancient exapted transposable elements promote nuclear enrichment of human long noncoding RNAs. Genome Res. 2019;29: 208–222. doi:10.1101/gr.229922.117

18. Denning DW. Global incidence and mortality of severe fungal disease. Lancet Infect Dis. 2024;24: e428–e438. doi:10.1016/S1473-3099(23)00692-8

19. Kujawa M. Some Naturally Occurring Substances: Food Items and Constituents, Heterocyclic Aromatic Amines and Mycotoxins. IARC Monographs on the Evaluation of Carcinogenic Risks to Humans, Vol. 56. Herausgegeben von der International Agency for Research on Cancer, World Health Organization. 599 Seiten, zahlr. Abb. und Tab. World Health Organization. Geneva 1993. Preis: 95, — Sw.fr; 95,50 US $. Food Nahr. 1994;38: 351–351. doi:10.1002/food.19940380335

20. Trail F, Mahanti N, Rarick M, Mehigh R, Liang SH, Zhou R, et al. Physical and transcriptional map of an aflatoxin gene cluster in Aspergillus parasiticus and functional disruption of a gene involved early in the aflatoxin pathway. Appl Environ Microbiol. 1995;61: 2665–2673. doi:10.1128/aem.61.7.2665-2673.1995

21. Cairns TC, Nai C, Meyer V. How a fungus shapes biotechnology: 100 years of Aspergillus niger research. Fungal Biol Biotechnol. 2018;5: 13. doi:10.1186/s40694-018-0054-5

22. Brandl J, Andersen MR. Aspergilli: Models for systems biology in filamentous fungi. Curr Opin Syst Biol. 2017;6: 67–73. doi:10.1016/j.coisb.2017.09.005

23. Steenwyk JL, Shen X-X, Lind AL, Goldman GH, Rokas A. A Robust Phylogenomic Time Tree for Biotechnologically and Medically Important Fungi in the Genera *Aspergillus* and *Penicillium*. Boyle JP, editor. mBio. 2019;10: e00925–19. doi:10.1128/mBio.00925-19

24. Vadlapudi V, Borah N, Yellusani KR, Gade S, Reddy P, Rajamanikyam M, et al. Aspergillus Secondary Metabolite Database, a resource to understand the Secondary metabolome of Aspergillus genus. Sci Rep. 2017;7: 7325. doi:10.1038/s41598-017-07436-w

25. Keller NP. Fungal secondary metabolism: regulation, function and drug discovery. Nat Rev Microbiol. 2019;17: 167–180. doi:10.1038/s41579-018-0121-1

26. Caceres I, Al Khoury A, El Khoury R, Lorber S. P. Oswald I, El Khoury A, et al. Aflatoxin Biosynthesis and Genetic Regulation: A Review. Toxins. 2020;12: 150. doi:10.3390/toxins12030150

27. Pfannenstiel BT, Keller NP. On top of biosynthetic gene clusters: How epigenetic machinery influences secondary metabolism in fungi. Biotechnol Adv. 2019;37: 107345. doi:10.1016/j.biotechadv.2019.02.001

28. Wang W, Yu Y, Keller NP, Wang P. Presence, Mode of Action, and Application of Pathway Specific Transcription Factors in Aspergillus Biosynthetic Gene Clusters. Int J Mol Sci. 2021;22: 8709. doi:10.3390/ijms22168709

29. Carriel CC, Halberg-Spencer SA, Kotvanova M, Pyne S, Park SC, Seo H, et al. A network-based model of *Aspergillus fumigatus* elucidates regulators of development and defensive natural products of an opportunistic pathogen. Nucleic Acids Res. 2026;54: gkaf1439. doi:10.1093/nar/gkaf1439

30. Kwon MJ, Steiniger C, Cairns TC, Wisecaver JH, Lind AL, Pohl C, et al. Beyond the Biosynthetic Gene Cluster Paradigm: Genome-Wide Coexpression Networks Connect Clustered and Unclustered Transcription Factors to Secondary Metabolic Pathways. Goldman GH, editor. Microbiol Spectr. 2021;9: e00898–21. doi:10.1128/Spectrum.00898-21

31. Staněk D. Long non-coding RNAs and splicing. Hon CC, editor. Essays Biochem. 2021;65: 723–729. doi:10.1042/EBC20200087

32. Kopp F, Mendell JT. Functional Classification and Experimental Dissection of Long Noncoding RNAs. Cell. 2018;172: 393–407. doi:10.1016/j.cell.2018.01.011

33. Godet A-C, Roussel E, Laugero N, Morfoisse F, Lacazette E, Garmy-Susini B, et al. Translational control by long non-coding RNAs. Biochimie. 2024;217: 42–53. doi:10.1016/j.biochi.2023.08.015

34. Cemel IA, Ha N, Schermann G, Yonekawa S, Brunner M. The coding and noncoding transcriptome of Neurospora crassa. BMC Genomics. 2017;18: 978. doi:10.1186/s12864-017-4360-8

35. Hovhannisyan H, Gabaldón T. The long non-coding RNA landscape of Candida yeast pathogens. Nat Commun. 2021;12: 7317. doi:10.1038/s41467-021-27635-4

36. Glad HM, Tralamazza SM, Croll D. The expression landscape and pangenome of long non-coding RNA in the fungal wheat pathogen Zymoseptoria tritici. Microb Genomics. 2023;9. doi:10.1099/mgen.0.001136

37. Kutter C, Watt S, Stefflova K, Wilson MD, Goncalves A, Ponting CP, et al. Rapid Turnover of Long Noncoding RNAs and the Evolution of Gene Expression. Bartel DP, editor. PLoS Genet. 2012;8: e1002841. doi:10.1371/journal.pgen.1002841

38. Pegueroles C, Iraola-Guzmán S, Chorostecki U, Ksiezopolska E, Saus E, Gabaldón T. Transcriptomic analyses reveal groups of co-expressed, syntenic lncRNAs in four species of the genus *Caenorhabditis*. RNA Biol. 2019;16: 320–329. doi:10.1080/15476286.2019.1572438

39. Fedorova ND, Khaldi N, Joardar VS, Maiti R, Amedeo P, Anderson MJ, et al. Genomic Islands in the Pathogenic Filamentous Fungus Aspergillus fumigatus. Richardson PM, editor. PLoS Genet. 2008;4: e1000046. doi:10.1371/journal.pgen.1000046

40. Galagan JE, Calvo SE, Cuomo C, Ma L-J, Wortman JR, Batzoglou S, et al. Sequencing of Aspergillus nidulans and comparative analysis with A. fumigatus and A. oryzae. Nature. 2005;438: 1105–1115. doi:10.1038/nature04341

41. Zhang X, Leahy I, Collemare J, Seidl MF. Genomic Localization Bias of Secondary Metabolite Gene Clusters and Association with Histone Modifications in *Aspergillus*. Giraud T, editor. Genome Biol Evol. 2024;16: evae228. doi:10.1093/gbe/evae228

42. Barber AE, Sae-Ong T, Kang K, Seelbinder B, Li J, Walther G, et al. Aspergillus fumigatus pan-genome analysis identifies genetic variants associated with human infection. Nat Microbiol. 2021;6: 1526–1536. doi:10.1038/s41564-021-00993-x

43. Langfelder P, Horvath S. WGCNA: an R package for weighted correlation network analysis. BMC Bioinformatics. 2008;9: 559. doi:10.1186/1471-2105-9-559

44. Zhang B, Horvath S. A General Framework for Weighted Gene Co-Expression Network Analysis. Stat Appl Genet Mol Biol. 2005;4. doi:10.2202/1544-6115.1128

45. Wiemann P, Perevitsky A, Lim FY, Shadkchan Y, Knox BP, Landero Figueora JA, et al. Aspergillus fumigatus Copper Export Machinery and Reactive Oxygen Intermediate Defense Counter Host Copper-Mediated Oxidative Antimicrobial Offense. Cell Rep. 2017;19: 1008–1021. doi:10.1016/j.celrep.2017.04.019

46. Schrettl M, Haas H. Iron homeostasis—Achilles’ heel of Aspergillus fumigatus? Curr Opin Microbiol. 2011;14: 400–405. doi:10.1016/j.mib.2011.06.002

47. Lanze CE, Gandra RM, Foderaro JE, Swenson KA, Douglas LM, Konopka JB. Plasma Membrane MCC/Eisosome Domains Promote Stress Resistance in Fungi. Microbiol Mol Biol Rev. 2020;84: e00063–19. doi:10.1128/MMBR.00063-19

48. Jong T, Stack CM, Moffitt MC, Morton CO. An Introduction to the Influence of Nutritional Factors on the Pathogenesis of Opportunist Fungal Pathogens in Humans. Pathogens. 2025;14: 335. doi:10.3390/pathogens14040335

49. Beauvais A, Latgé J-P. *Aspergillus* Biofilm *In Vitro* and *In Vivo*. Ghannoum M, Parsek M, Whiteley M, Mukherjee P, editors. Microbiol Spectr. 2015;3: 3.4.08. doi:10.1128/microbiolspec.MB-0017-2015

50. Puerner C, Morelli KA, Kerkaert JD, Jones JT, Quinn KG, DeMichaelis N, et al. Transcriptional and metabolic modeling analyses of developing *Aspergillus fumigatus* biofilms reveal metabolic shifts required for biofilm maturation. Zhai B, editor. mSphere. 2025;10: e00752–25. doi:10.1128/msphere.00752-25

51. Baril T, Croll D. MycoMobilome: A community-focused non-redundant database of transposable element consensus sequences for the fungal kingdom. Genomics; 2025. doi:10.1101/2025.10.28.685023

52. Gladyshev E. Repeat-Induced Point Mutation and Other Genome Defense Mechanisms in Fungi. Heitman J, Stukenbrock EH, editors. Microbiol Spectr. 2017;5: 5.4.02. doi:10.1128/microbiolspec.FUNK-0042-2017

53. Wang L, Sun Y, Sun X, Yu L, Xue L, He Z, et al. Repeat-induced point mutation in Neurospora crassa causes the highest known mutation rate and mutational burden of any cellular life. Genome Biol. 2020;21: 142. doi:10.1186/s13059-020-02060-w

54. Hane JK, Oliver RP. In silico reversal of repeat-induced point mutation (RIP) identifies the origins of repeat families and uncovers obscured duplicated genes. BMC Genomics. 2010;11: 655. doi:10.1186/1471-2164-11-655

55. Palmer JM, Keller NP. Secondary metabolism in fungi: does chromosomal location matter? Curr Opin Microbiol. 2010;13: 431–436. doi:10.1016/j.mib.2010.04.008

56. Badet T, Croll D. Phylogenomic signatures of repeat-induced point mutations across the fungal kingdom. Kelleher ES, editor. PLOS Biol. 2025;23: e3003433. doi:10.1371/journal.pbio.3003433

57. Camilleri-Robles C, Amador R, Klein CC, Guigó R, Corominas M, Ruiz-Romero M. Genomic and functional conservation of lncRNAs: lessons from flies. Mamm Genome. 2022;33: 328–342. doi:10.1007/s00335-021-09939-4

58. Engreitz JM, Haines JE, Perez EM, Munson G, Chen J, Kane M, et al. Local regulation of gene expression by lncRNA promoters, transcription and splicing. Nature. 2016;539: 452–455. doi:10.1038/nature20149

59. Gil N, Ulitsky I. Regulation of gene expression by cis-acting long non-coding RNAs. Nat Rev Genet. 2020;21: 102–117. doi:10.1038/s41576-019-0184-5

60. Ransohoff JD, Wei Y, Khavari PA. The functions and unique features of long intergenic non-coding RNA. Nat Rev Mol Cell Biol. 2018;19: 143–157. doi:10.1038/nrm.2017.104

61. Hatmaker EA, Barber AE, Drott MT, Sauters TJC, Gumilang A, Alastruey-Izquierdo A, et al. Population structure in a fungal human pathogen is potentially linked to pathogenicity. Nat Commun. 2025;16: 7594. doi:10.1038/s41467-025-62777-9

62. Bok JW, Keller NP. LaeA, a Regulator of Secondary Metabolism in *Aspergillus* spp. Eukaryot Cell. 2004;3: 527–535. doi:10.1128/EC.3.2.527-535.2004

63. Kale SP, Milde L, Trapp MK, Frisvad JC, Keller NP, Bok JW. Requirement of LaeA for secondary metabolism and sclerotial production in Aspergillus flavus. Fungal Genet Biol. 2008;45: 1422–1429. doi:10.1016/j.fgb.2008.06.009

64. Mosquera S, Ginésy M, Bocos-Asenjo IT, Amin H, Diez-Hermano S, Diez JJ, et al. Spray-induced gene silencing to control plant pathogenic fungi: A step-by-step guide. J Integr Plant Biol. 2025;67: 801–825. doi:10.1111/jipb.13848

65. Ghosh S, Patra S, Ray S. A Combinatorial Nanobased Spray-Induced Gene Silencing Technique for Crop Protection and Improvement. ACS Omega. 2023;8: 22345–22351. doi:10.1021/acsomega.3c01968

66. Li N, Xu X, Li J, Hull JJ, Chen L, Liang G. A spray-induced gene silencing strategy for Spodoptera frugiperda oviposition inhibition using nanomaterial-encapsulated dsEcR. Int J Biol Macromol. 2024;281: 136503. doi:10.1016/j.ijbiomac.2024.136503

67. Gilbert MK, Majumdar R, Rajasekaran K, Chen Z-Y, Wei Q, Sickler CM, et al. RNA interference-based silencing of the alpha-amylase (amy1) gene in Aspergillus flavus decreases fungal growth and aflatoxin production in maize kernels. Planta. 2018;247: 1465–1473. doi:10.1007/s00425-018-2875-0

68. Thakare D, Zhang J, Wing RA, Cotty PJ, Schmidt MA. Aflatoxin-free transgenic maize using host-induced gene silencing. Sci Adv. 2017;3: e1602382. doi:10.1126/sciadv.1602382

69. Zhuang Z, Sun M, Wu D, Ma D, Chen L, Pan X, et al. COMPASS subunit Bre2 regulates chromatin remodeler Arp9 to control Aspergillus flavus aflatoxin synthesis and virulence. Nat Commun. 2026;17: 1862. doi:10.1038/s41467-026-69877-0

70. Ma D, Yao Y, Yang C, Lin H, Sun M, Gao Y, et al. The chromatin remodeling factor Arp9 modulates drug-resistance and plays a key role in aflatoxins biosynthesis under mammalian-physiological-temperature in Aspergillus flavus. Liu H, editor. PLOS Pathog. 2026;22: e1014021. doi:10.1371/journal.ppat.1014021

71. Gervais NC, Rogers RKJ, Robin MR, Shapiro RS. HyperdCas12a-based multiplexed genetic regulation in *Candida albicans*. Nucleic Acids Res. 2025;53: gkaf1402. doi:10.1093/nar/gkaf1402

72. Alvarez-Jarreta J, Amos B, Aurrecoechea C, Bah S, Barba M, Barreto A, et al. VEuPathDB: the eukaryotic pathogen, vector and host bioinformatics resource center in 2023. Nucleic Acids Res. 2024;52: D808–D816. doi:10.1093/nar/gkad1003

73. Basenko EY, Pulman JA, Shanmugasundram A, Harb OS, Crouch K, Starns D, et al. FungiDB: An Integrated Bioinformatic Resource for Fungi and Oomycetes. J Fungi. 2018;4: 39. doi:10.3390/jof4010039

74. Blin K, Shaw S, Augustijn HE, Reitz ZL, Biermann F, Alanjary M, et al. antiSMASH 7.0: new and improved predictions for detection, regulation, chemical structures and visualisation. Nucleic Acids Res. 2023;51: W46–W50. doi:10.1093/nar/gkad344

75. Jones P, Binns D, Chang H-Y, Fraser M, Li W, McAnulla C, et al. InterProScan 5: genome-scale protein function classification. Bioinformatics. 2014;30: 1236–1240. doi:10.1093/bioinformatics/btu031

76. Patro R, Duggal G, Love MI, Irizarry RA, Kingsford C. Salmon provides fast and bias-aware quantification of transcript expression. Nat Methods. 2017;14: 417–419. doi:10.1038/nmeth.4197

77. Snakemake—a scalable bioinformatics workflow engine | Bioinformatics | Oxford Academic. [cited 19 May 2026]. Available: https://academic.oup.com/bioinformatics/article/28/19/2520/290322

78. Ewels P, Magnusson M, Lundin S, Käller M. MultiQC: summarize analysis results for multiple tools and samples in a single report. Bioinformatics. 2016;32: 3047–3048. doi:10.1093/bioinformatics/btw354

79. Chen S, Zhou Y, Chen Y, Gu J. fastp: an ultra-fast all-in-one FASTQ preprocessor. Bioinformatics. 2018;34: i884–i890. doi:10.1093/bioinformatics/bty560

80. Dobin A, Davis CA, Schlesinger F, Drenkow J, Zaleski C, Jha S, et al. STAR: ultrafast universal RNA-seq aligner. Bioinformatics. 2013;29: 15–21. doi:10.1093/bioinformatics/bts635

81. Pertea M, Pertea GM, Antonescu CM, Chang T-C, Mendell JT, Salzberg SL. StringTie enables improved reconstruction of a transcriptome from RNA-seq reads. Nat Biotechnol. 2015;33: 290–295. doi:10.1038/nbt.3122

82. Pertea G, Pertea M. GFF Utilities: GffRead and GffCompare. F1000Research. 2020;9: 304. doi:10.12688/f1000research.23297.2

83. Cribbs AP, Luna-Valero S, George C, Sudbery IM, Berlanga-Taylor AJ, Sansom SN, et al. CGAT-core: a python framework for building scalable, reproducible computational biology workflows. F1000Research. 2019;8: 377. doi:10.12688/f1000research.18674.1

84. Kang Y-J, Yang D-C, Kong L, Hou M, Meng Y-Q, Wei L, et al. CPC2: a fast and accurate coding potential calculator based on sequence intrinsic features. Nucleic Acids Res. 2017;45: W12–W16. doi:10.1093/nar/gkx428

85. Wucher V, Legeai F, Hédan B, Rizk G, Lagoutte L, Leeb T, et al. FEELnc: a tool for long non-coding RNA annotation and its application to the dog transcriptome. Nucleic Acids Res. 2017; gkw1306. doi:10.1093/nar/gkw1306

86. McGinnis S, Madden TL. BLAST: at the core of a powerful and diverse set of sequence analysis tools. Nucleic Acids Res. 2004;32: W20–W25. doi:10.1093/nar/gkh435

87. Nawrocki EP, Kolbe DL, Eddy SR. Infernal 1.0: inference of RNA alignments. Bioinformatics. 2009;25: 1335–1337. doi:10.1093/bioinformatics/btp157

88. Kalvari I, Nawrocki EP, Argasinska J, Quinones-Olvera N, Finn RD, Bateman A, et al. Non-Coding RNA Analysis Using the Rfam Database. Curr Protoc Bioinforma. 2018;62: e51. doi:10.1002/cpbi.51

89. Jacques Dainat, Darío Hereñú, Dr. K. D. Murray, Ed Davis, Ivan Ugrin, Kathryn Crouch, et al. NBISweden/AGAT: AGAT-v1.4.1. Zenodo; 2024. doi:10.5281/ZENODO.3552717

90. Lovell JT, Sreedasyam A, Schranz ME, Wilson M, Carlson JW, Harkess A, et al. GENESPACE tracks regions of interest and gene copy number variation across multiple genomes. eLife. 2022;11: e78526. doi:10.7554/eLife.78526

91. Emms DM, Kelly S. OrthoFinder: phylogenetic orthology inference for comparative genomics. Genome Biol. 2019;20: 238. doi:10.1186/s13059-019-1832-y

92. Quinlan AR, Hall IM. BEDTools: a flexible suite of utilities for comparing genomic features. Bioinformatics. 2010;26: 841–842. doi:10.1093/bioinformatics/btq033

93. Liao Y, Smyth GK, Shi W. featureCounts: an efficient general purpose program for assigning sequence reads to genomic features. Bioinformatics. 2014;30: 923–930. doi:10.1093/bioinformatics/btt656

94. Baril T, Galbraith J, Hayward A. Earl Grey: A Fully Automated User-Friendly Transposable Element Annotation and Analysis Pipeline. Arkhipova I, editor. Mol Biol Evol. 2024;41: msae068. doi:10.1093/molbev/msae068

95. Lawrence M, Huber W, Pagès H, Aboyoun P, Carlson M, Gentleman R, et al. Software for Computing and Annotating Genomic Ranges. Prlic A, editor. PLoS Comput Biol. 2013;9: e1003118. doi:10.1371/journal.pcbi.1003118

96. Pedersen TL. tidygraph: A Tidy API for Graph Manipulation. 2017. p. 1.3.1. doi:10.32614/CRAN.package.tidygraph

97. Pedersen TL. ggraph: An Implementation of Grammar of Graphics for Graphs and Networks. 2017. p. 2.2.1. doi:10.32614/CRAN.package.ggraph

98. Xu S, Hu E, Cai Y, Xie Z, Luo X, Zhan L, et al. Using clusterProfiler to characterize multiomics data. Nat Protoc. 2024;19: 3292–3320. doi:10.1038/s41596-024-01020-z

99. Love MI, Huber W, Anders S. Moderated estimation of fold change and dispersion for RNA-seq data with DESeq2. Genome Biol. 2014;15: 550. doi:10.1186/s13059-014-0550-8

100. Frawley D, Greco C, Oakley B, Alhussain MM, Fleming AB, Keller NP, et al. The tetrameric pheromone module SteC-MkkB-MpkB-SteD regulates asexual sporulation, sclerotia formation and aflatoxin production in Aspergillus flavus. Cell Microbiol. 2020;22: e13192. doi:10.1111/cmi.13192

101. Bok JW, Soukup AA, Chadwick E, Chiang Y-M, Wang CCC, Keller NP. VeA and MvlA repression of the cryptic orsellinic acid gene cluster in Aspergillus nidulans involves histone 3 acetylation. Mol Microbiol. 2013;89: 963–974. doi:10.1111/mmi.12326

102. Green MR, Sambrook J. Molecular cloning: a laboratory manual. 4th ed. Cold Spring Harbor: Cold Spring Harbor laboratory press; 2012.

103. Álvarez-Escribano I, Sasse C, Bok JW, Na H, Amirebrahimi M, Lipzen A, et al. Genome sequencing of evolved aspergilli populations reveals robust genomes, transversions in A. flavus, and sexual aberrancy in non-homologous end-joining mutants. BMC Biol. 2019;17: 88. doi:10.1186/s12915-019-0702-0

104. Barbas C, Dams A, Majors RE. Separation of aflatoxins by HPLC. Agilent Technologies; 2005. Available: https://www.agilent.com/Library/applications/5989-3634EN.pdf

105. Wickham H, Chang W, Henry L, Pedersen TL, Takahashi K, Wilke C, et al. ggplot2: Create Elegant Data Visualisations Using the Grammar of Graphics. 2007. p. 3.5.2. doi:10.32614/CRAN.package.ggplot2

106. Wilke CO. ggridges: Ridgeline Plots in “ggplot2.” 2017. p. 0.5.6. doi:10.32614/CRAN.package.ggridges

107. Kassambara A. ggpubr: “ggplot2” Based Publication Ready Plots. 2016. p. 0.6.0. doi:10.32614/CRAN.package.ggpubr

