## Supplementary Information for "An *Aspergillus* lncRNA atlas reveals a novel modulator of aflatoxin biosynthesis"

### Table of content

A. **Supplementary Figures**

- Supplementary Figure S1: Structural and genomic organization of lncRNAs across *Aspergillus* species
- Supplementary Figure S2: Subtelomeric localization influences TE content of Aspergillus intergenic lncRNAs.
- Supplementary Figure S3: Network topology analysis for soft-thresholding power selection
- Supplementary Figure S4: Topological architecture and composition of *Aspergillus* co-expression networks
- Supplementary Figure S5: Module preservation analysis of the *A. fumigatus* co-expression network
- Supplementary Figure S6: Identification and characterization of the lincRNA *aflalinc* in *A. flavus*

B. **Supplementary Tables**

- Supplementary Table S1: List of SRA projects analyzed in this study
- Supplementary Table S2: Comparative genomics and Chi-square statistics for subtelomeric enrichment of lncRNAs
- Supplementary Table S3: Syntenic families of intergenic lncRNAs across *Aspergillus* species
- Supplementary Table S4: Comprehensive summary of transposable elements (TEs)
- Supplementary Table S5: *Aspergillus* lncRNA functional module annotations, including:
  - Table I: Species-wide network statistics
  - Table II: High confidence lncRNA co-expression modules
  - Table III: Aflatoxin-related module in *A. flavus*
  - Table IV: Functional categorization of modules
  - Table V: Complete module list
  - Table VI: Manually curated functional categories of lncRNA-enriched modules
- Supplementary Table S6: LincRNAs with biofilm-stage classifications
- Supplementary Table S7: Details of *aflalinc* experimental datasets
- Supplementary Table S8: Spearman correlation coefficients (ρ) for lincRNA-PCGs pairs within 15 kb genomic proximity.

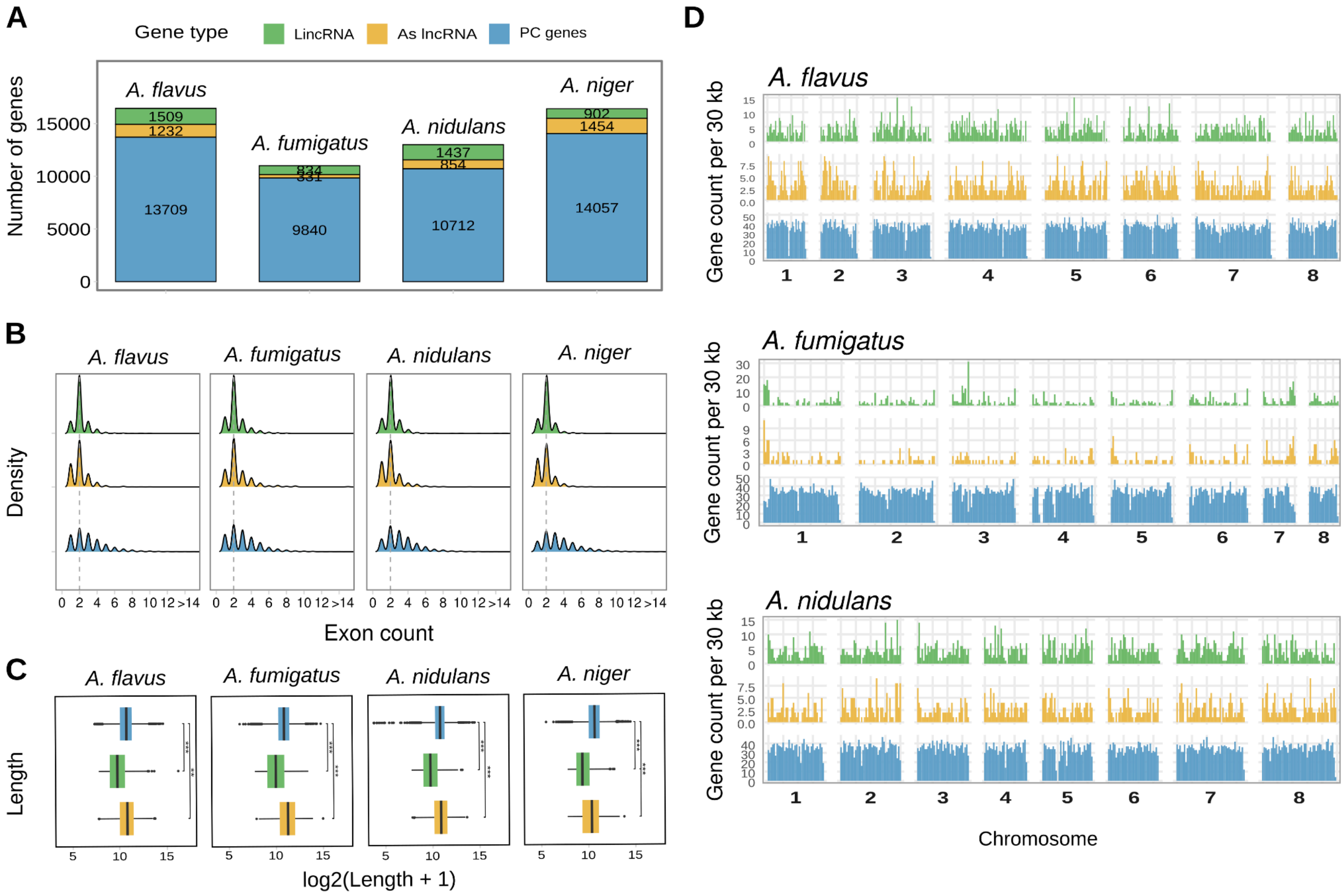


**Supplementary Figure S1. Structural and genomic organization of lncRNAs across *Aspergillus* species**. (A) Stack plots illustrating the total annotated gene counts for *A. flavus, A. fumigatus, A. nidulans,* and *A. niger.* The lncRNA annotations represent the newly generated non-coding gene models produced in this study, while PGC counts represent those in the reference annotation. (B) Density distributions for lncRNAs vs. mRNAs. (C) Lengths of proteins, lincRNAs, and As lncRNAs. Length values are log2-transformed for clarity. Statistical significance was assessed using the Wilcoxon rank sum test with Benjamini-Hochberg correction; *p < 0.05, **p < 0.01, ***p < 0.001. (D) Chromosomal distribution of annotated gene types across *Aspergillus* genomes. Each genomic feature is colored according to its annotation class: protein-coding genes (blue), As lncRNAs (orange), and lincRNAs (green).

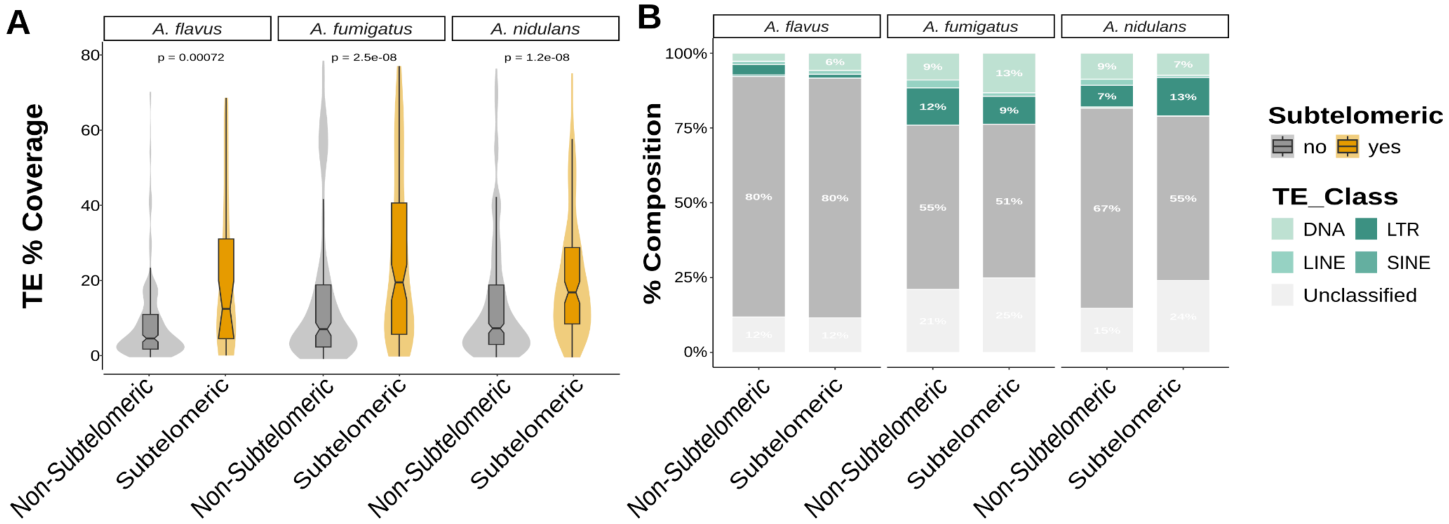


**Supplementary Figure S2. Subtelomeric localization influences TE content of *Aspergillus* intergenic lncRNAs.** (A) Distribution of TE sequence coverage in subtelomeric versus non-subtelomeric lincRNAs across three *Aspergillus* species. Values represent the percentage of exon bases that overlap annotated TE sequence. Only TE-associated lincRNAs are shown. Significance was assessed using Wilcoxon rank-sum tests with Bonferroni correction. (B) Composition of TE classes among TE-associated lincRNAs. *A. niger* is excluded from both panels due to its scaffold-level genome assembly precluding reliable subtelomere annotation.

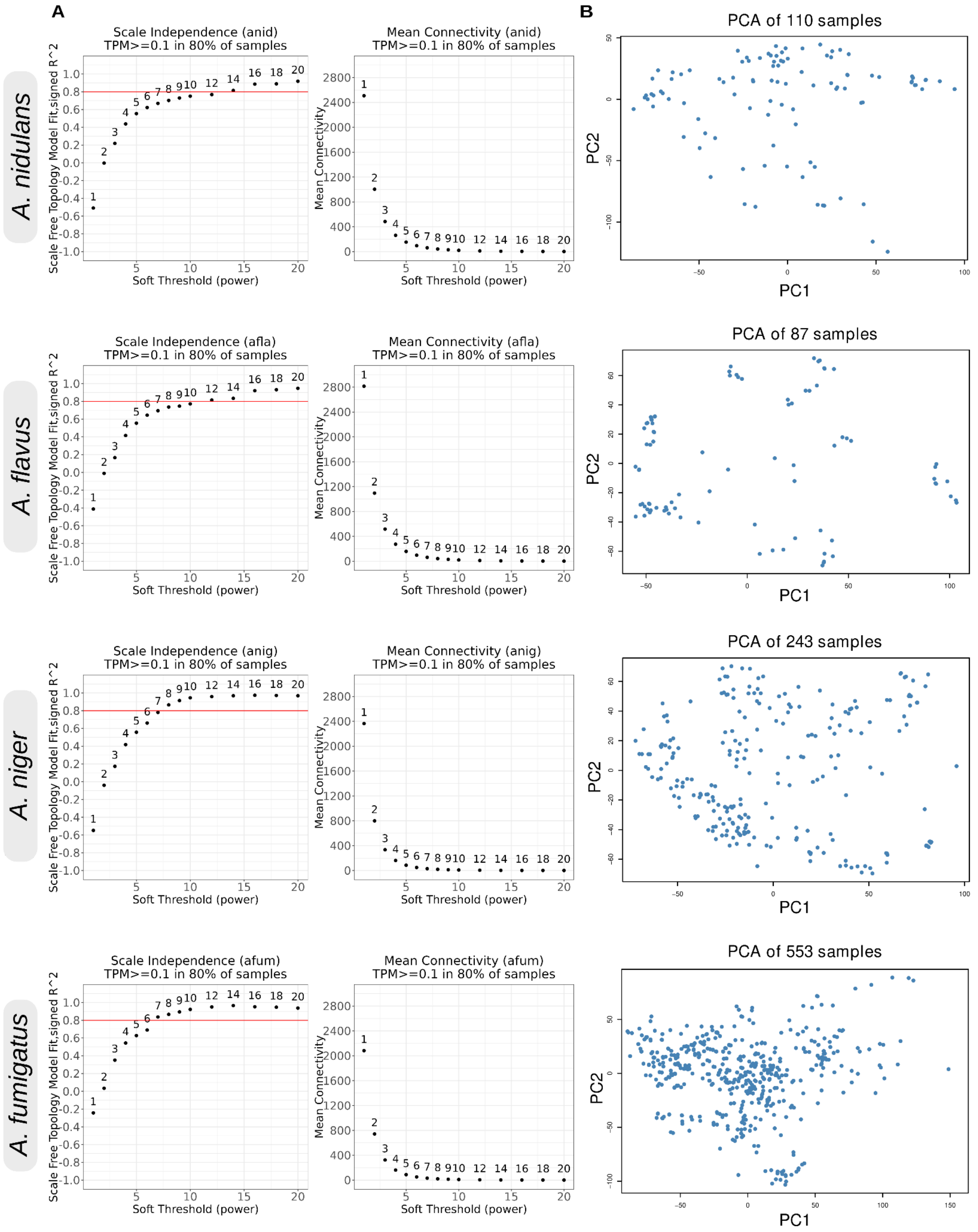


**Supplementary Figure S3. Network topology analysis for soft-thresholding power selection.** (A) Analysis of the scale-free fit index ($R^2$) for various soft-thresholding powers and the mean connectivity as a function of the soft-thresholding power. The horizontal line indicates the $R^2$ = 0.80 threshold used for network construction. (B) Principal component analysis (PCA) of all samples used for network construction.

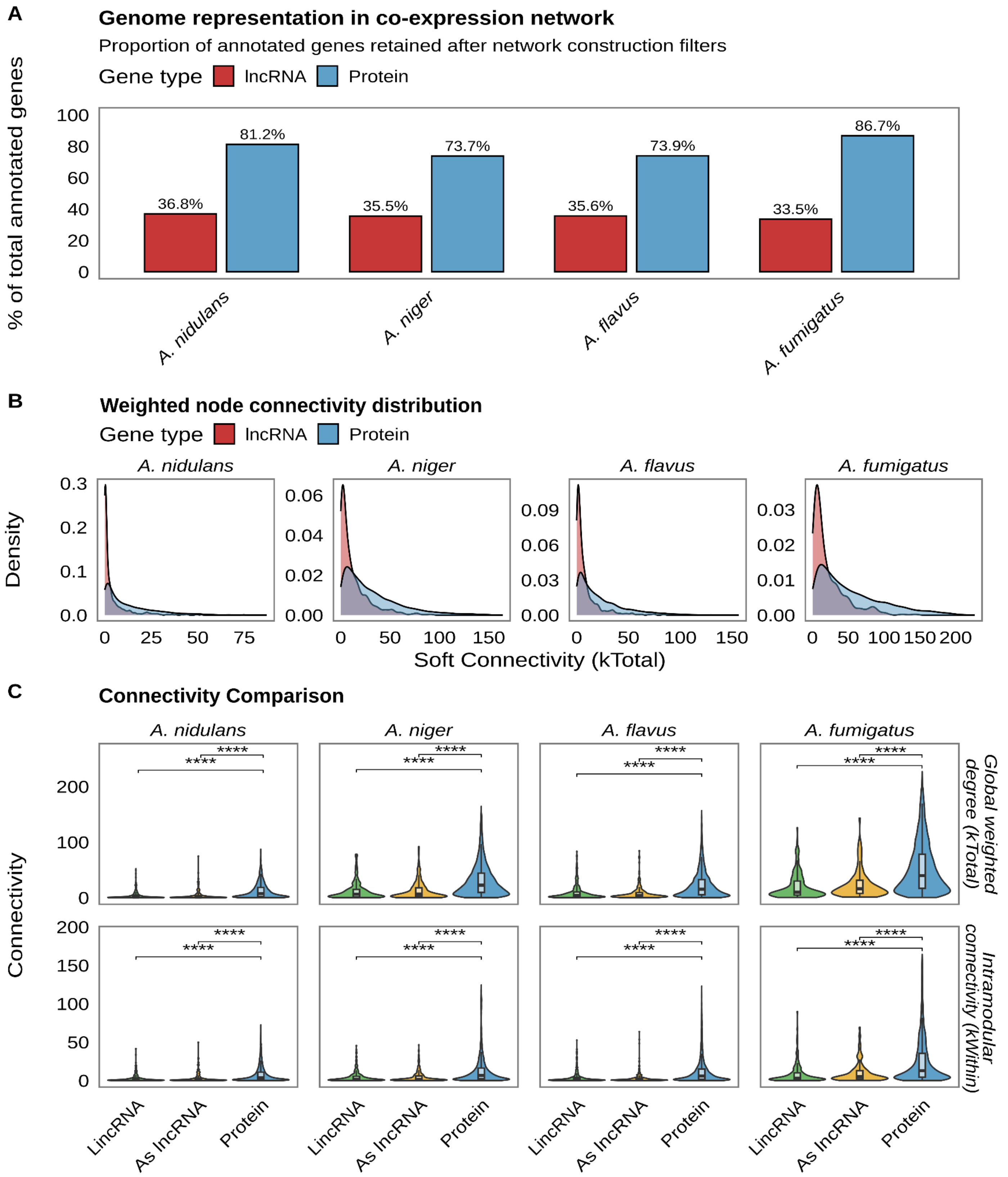


**Supplementary Figure S4. Co-expression network properties of lncRNAs versus protein-coding genes in *Aspergillus*.** (A) Bar plots showing the proportion of protein-coding genes (blue) and lncRNAs (red) that were retained in the final co-expression network after filtering for expression TPM ≥ 0.1 in at least 80% of samples. (B) Kernel density plots showing the distribution of global connectivity ($k_{Total}$) for lncRNAs (red) and protein-coding genes (blue) in the four *Aspergillus* species. Connectivity represents the sum of adjacency weights. (C) Violin and boxplots illustrating the distribution of global connectivity ($k_{Total}$) and intramodular connectivity ($k_{Within}$) for protein-coding genes, As lncRNA, and lincRNAs. Significance was assessed using two-sided Wilcoxon rank-sum tests with Benjamini-Hochberg correction; ****p < 0.0001, ***p < 0.0001, **p < 0.001, *p < 0.05.

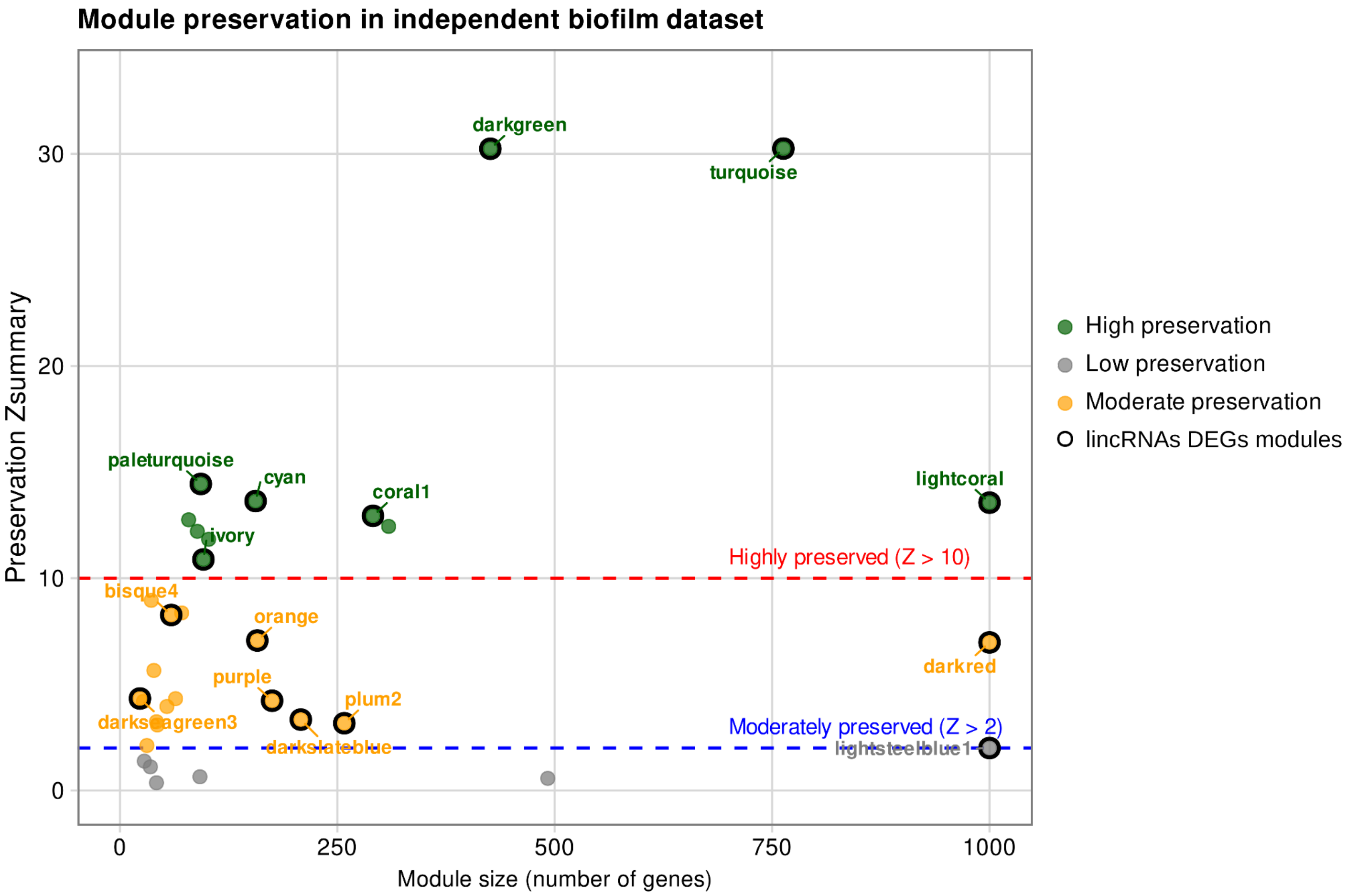


**Supplementary Figure S5. Module preservation analysis confirms network stability across an independent dataset.** Scatter plot of module size vs. preservation Zsummary (Z) statistics for all modules defined in the global *A. fumigatus* co-expression network when tested against an independent biofilm maturation dataset. Each point represents a module, colored by preservation category: high preservation ($Z > 10$, green), moderate preservation ($2 < Z < 10$, orange), and low preservation ($Z < 2$, grey). Dashed red and blue lines indicate $Z = 10$ and $Z = 2$ thresholds, respectively. Modules containing differentially expressed lincRNAs are highlighted with black circles and labeled by name.

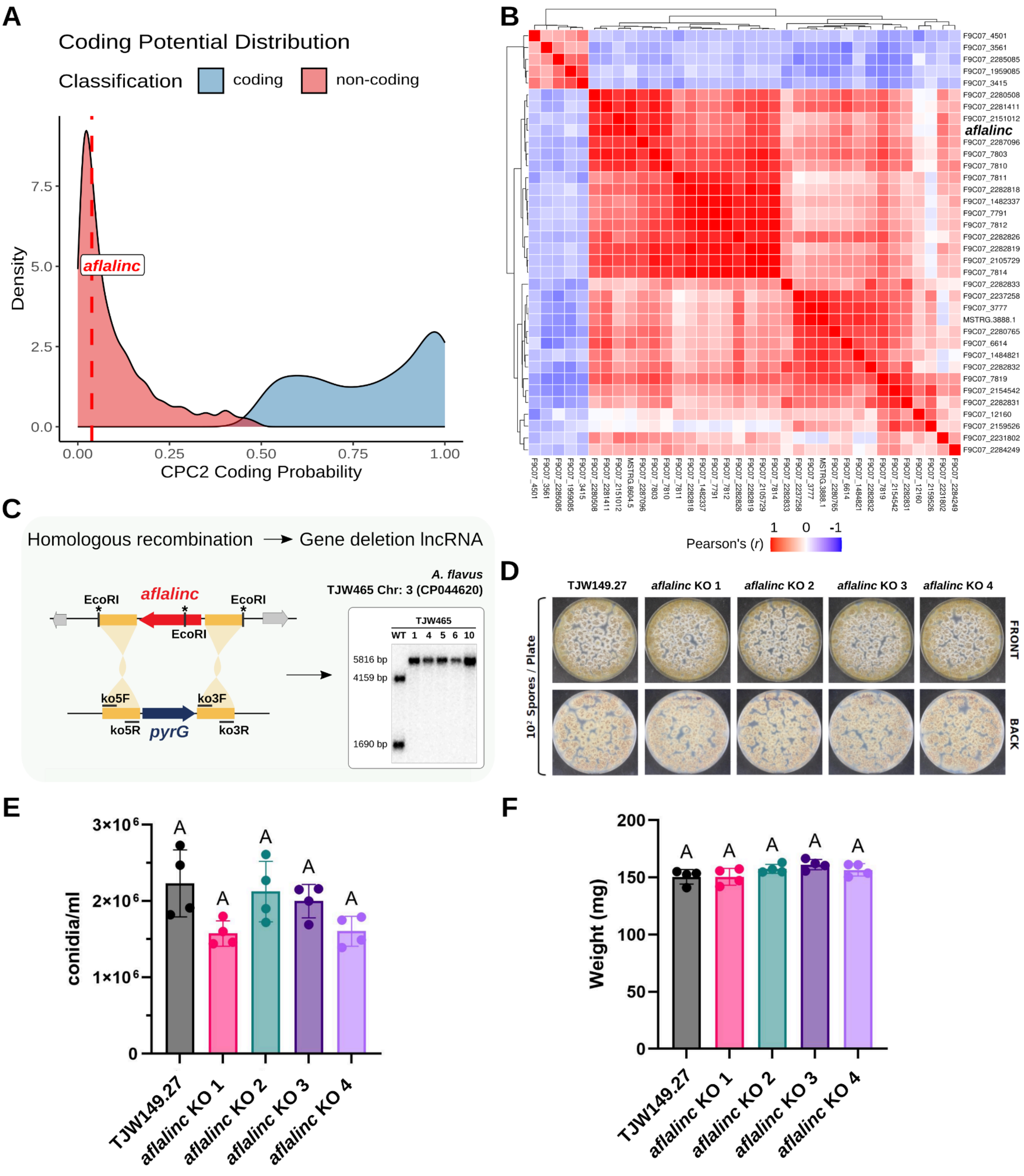


**Supplementary Figure S6. Identification and characterization of the intergenic lncRNA *aflalinc*.** (A) Density distribution of coding probability scores for PGCs (blue) and lncRNAs (red) in *A. flavus*. The dashed red line indicates the coding probability of *aflalinc* (score: 0.038), placing it firmly within the non-coding population. (B) Pearson correlation matrix of the salmon1 module containing *aflalinc* and many members of the aflatoxin BGC. (C) Schematic representation of the homologous recombination strategy used for deletion of the native *aflalinc* locus and replacement with the *pyrG* selectable marker. Southern blot analysis of genomic DNA confirming successful integration and the absence of ectopic insertions in four independent mutant strains (Δ*aflalinc* KO 1,2,3, and 4). (D) Representative images (front and back views) of colony morphology for the wild-type (WT) and four independent Δ*aflalinc* mutants after growth on glucose minimal medium at 30°C in the dark for 1 week, following inoculation with $10^2$ spores per plate. (E-F) Quantitative analysis of asexual development (E) and sclerotia biomass (F) in WT and Δ*aflalinc* mutants. Bars represent the mean ±SD of four biological replicates. Significance was determined using one-way ANOVA with Tukey's multiple-comparisons test in GraphPad Prism v10.6.1; letters indicate significant differences ($p < 0.05$). Values sharing the same letter are not significantly different.
